# Olfactory projection neuron rewiring in the brain of an ecological specialist

**DOI:** 10.1101/2024.07.26.605288

**Authors:** Benedikt R. Dürr, Enrico Bertolini, Suguru Takagi, Justine Pascual, Liliane Abuin, Giovanna Lucarelli, Richard Benton, Thomas O. Auer

**Author notes:** Corresponding author: Thomas O. Auer. equal contribution (Enrico Bertolini, Suguru Takagi, Justine Pascual).

## Abstract

Animals’ behaviours can vary greatly between even closely-related species. While changes in the sensory periphery have frequently been linked to species-specific behaviours, very little is known about if and how individual cell types in the central brain evolve. Here, we develop a set of advanced genetic tools to compare homologous neurons in *Drosophila sechellia* – which specialises on a single fruit – and *Drosophila melanogaster*. Through systematic morphological analysis of olfactory projection neurons (PNs), we reveal that global anatomy of these second-order neurons is conserved. However, high-resolution, quantitative comparisons identify a striking case of convergent rewiring of PNs in two distinct olfactory pathways critical for *D. sechellia*’s host location. Calcium imaging and labelling of pre-synaptic sites in these evolved PNs demonstrate that novel functional connections with third-order partners are formed in *D. sechellia*. This work demonstrates that peripheral sensory evolution is accompanied by highly-selective wiring changes in the central brain to facilitate ecological specialisation, and paves the way for systematic comparison of other cell types throughout the nervous system.

## Introduction

To prosper in defined ecological niches, animals exhibit specific behavioural programmes matching the requirements of their environment. Underlying these behaviours are neural circuits that receive external stimuli, integrate internal state and experience and control motor outputs. While behaviourally relevant circuits have been studied intensely in selected model species, the use of non-model organisms in comparative neuroscience is of growing interest^1–3^. Technological advances in gene editing and -omics methods facilitate the detailed analysis of multiple species to address how behaviours evolve through the alteration of genes, molecules, cells, and circuits^4,5^. Such comparisons help to detect hotspots of evolutionary change and decipher common schemes of brain and circuit evolution^3^. At the macroscopic level, the study of brain structures or number of cell types across large evolutionary time scales, has uncovered examples of brain re-organisation and divergence of cell numbers and types^6,7^. At shorter evolutionary distances, brain circuits modifications across species through altered gene expression^8^ or cell type composition^9^, among other changes, are likely to be more subtle. However, comparing closely related species may provide insights into recent changes that underlie behavioural divergence and help to understand how the accumulation of small alterations can eventually impact circuit output. Linking changes in neural circuits to species’ differences in behavioural output has, however, been technically challenging.

Sensory systems, at the interface of the brain and the environment, have been shown to be particularly prone to evolutionary adaptations^10^ and have provided the few rare examples where anatomical insights at the single cell level have been gained. For example, comparative connectomics after volumetric electron-microscopy imaging has pinpointed anatomical changes in defined microcircuits in the vertebrate retina that impact signal processing^11^. Synaptic rewiring in feeding circuits of two nematodes is linked to divergent predatory behaviours^12,13^. Similarly, in larval drosophilids, increased synaptic connections from the sensory periphery to shared downstream neurons explains species-specific sensitivity of nociceptive circuits^14^. Though these limited examples demonstrate the promise of comparative connectomics, this approach is currently limited to small volumes and very small sample sizes.

The olfactory system in flies, with its well delineated neural pathways and ability to label individual cell types is ideally suited to address the question of how individual circuits differ between species. This sensory system has a well-characterised, stereotypic neuroanatomy in *D. melanogaster*^15,16^: Olfactory information is sensed by olfactory sensory neurons (OSNs) located mainly on the third segment of the antenna. OSNs project their axons into the antennal lobe (AL) where they ramify in distinct olfactory glomeruli and connect to projection neurons (PNs) and local interneurons. PNs then transmit olfactory information to higher brain areas including the mushroom body (a centre for learning and memory) and the lateral horn (involved in innate behaviours)^17^.

The highly stereotyped nature of OSN-PN circuits has helped to uncover basic principles of neuronal differentiation, circuit wiring, and processing of olfactory information from the periphery to the central brain in *D. melanogaster*. PN dendritic and axonal growth is orchestrated by transcriptional programs which lead to coordinated development of AL connections between OSNs and downstream circuits^18–24^. PNs from individual glomeruli project to distinct zones within the LH in a highly stereotypic manner across individuals. This suggests that LH targeting is hardwired and leads to a functional subdivision of the LH neuropil.

This stereotypic anatomy has a direct impact on processing of olfactory information. Behaviourally relevant odorant valence is encoded by an odour-specific activation pattern of glomeruli in the AL^25^ – ranging from single glomeruli to weighted summation of multiple pathways. After pre-processing at the OSN-PN synapse and integration of local interneuron inputs, information is transmitted by PNs to the MB and LH where hedonic valences and odour intensities are represented as spatially segregated functional maps^26^. This spatial segregation helps to categorise odours based on their behavioural value^27,28^ and favours conversion of inputs onto downstream circuits with a defined behavioural role.

To understand how olfactory circuits evolve in response to changing environmental needs, *D. sechellia* is of particular interest: it is extremely specialized to life on a single host fruit *Morinda citrifolia* (“noni“) compared to the closely-related generalists *D. melanogaster* and *D. simulans* (**Fig. 1a**). Specific amino-acid changes in olfactory receptors^29–32^ increase *D. sechellia*’s sensitivity towards host odours and are accompanied by increases in OSN number^33^ which impact OSN-PN temporal response dynamics^33^. Despite the advances in understanding of peripheral olfactory evolution, we have very limited knowledge about relevant changes that have occurred in central olfactory pathways^29,34^. Laborious dye filling and photoactivation experiments^34^ suggest that there have been alterations in mushroom body connectivity across species. However, to gain single cell resolution in these experiments is very challenging.

**Figure 1.**
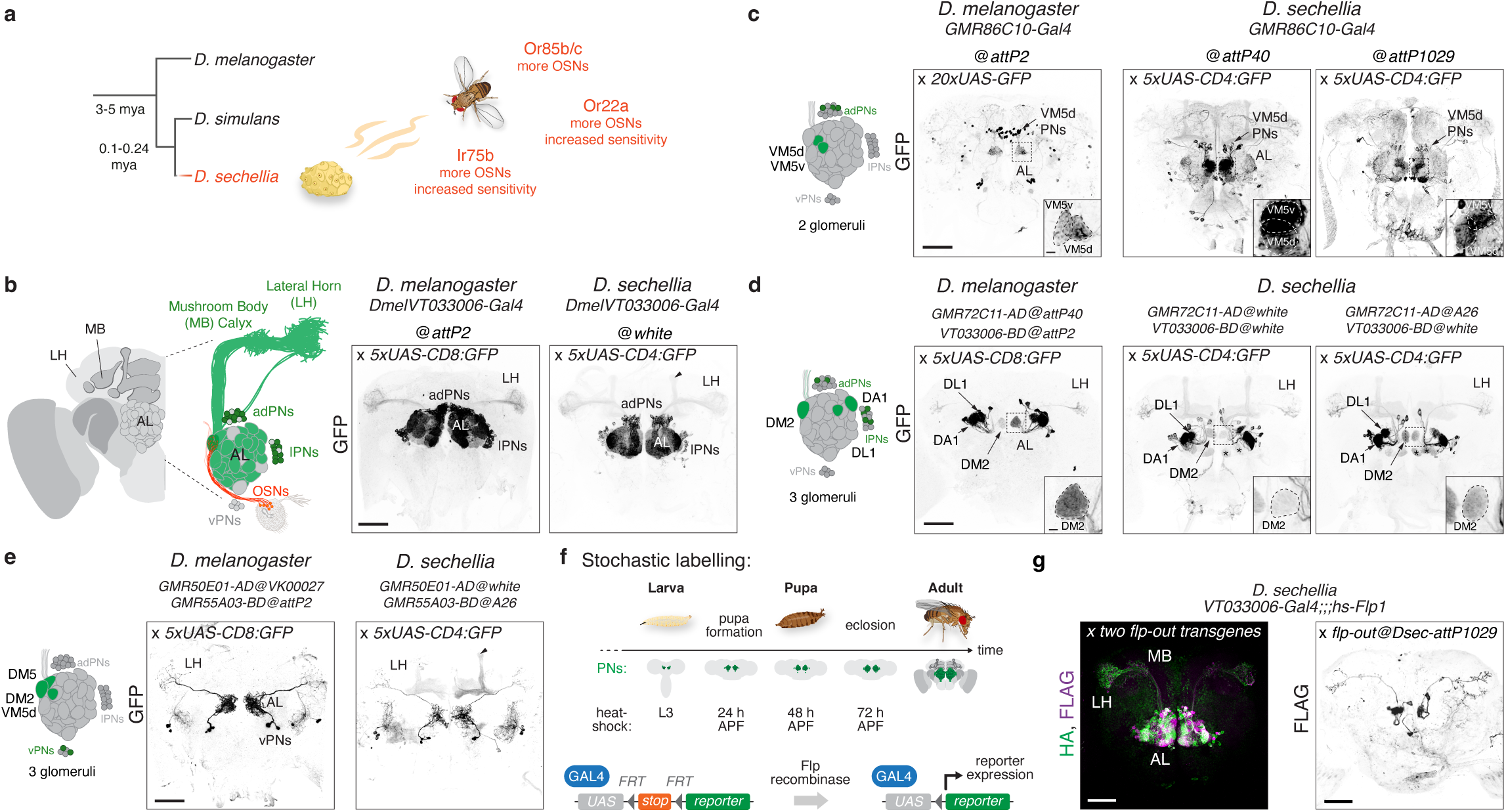
Generation of PN labelling tools in *D. sechellia*. **a)** Phylogenetic relationship of *D. melanogaster*, *D. simulans* and *D. sechellia*. The first two are cosmopolitan generalists while *D. sechellia* is an island endemic specialised on *Morinda citrifolia* (“noni”) fruit and shows several changes in its peripheral olfactory system. The two olfactory pathways expressing receptors Or85b/c and Or22a are essential for long-range attractive behaviour to noni in *D. sechellia*, the Ir75b pathway is involved in egg laying^29,58^. All three show an increase in OSN numbers in *D. sechellia*^33^ and the Ir75b and Or22a receptors increased sensitivity to host volatiles^29,31^. **b)** Left: Schematic of the fly brain and the olfactory system. Olfactory sensory neurons (OSNs, in orange) housed in sensilla on (mainly) the third antennal segment send their axons to glomerular structures in the antennal lobe (AL). Axons of neurons expressing the same receptor converge on the same glomerulus. Second order PNs with cell bodies in three clusters (anterior-dorsal, ad; lateral, l; ventral, v) around the AL connect via their dendrites to OSN axons and project axons to the mushroom body (MB) calyx and the lateral horn (LH). Right: Anti-GFP immunohistochemical staining of the *D. melanogaster* and *D. sechellia* central brain. The *VT033006-Gal4* transgenic line labels ad- and lPNs innervating 44 glomeruli^24^. Here and in other figures (if not indicated otherwise) stained brains were registered on a female reference brain of the respective species (*D. melanogaster:* IS2*, D. sechellia:* DsecF)^29,90^. Black arrowhead: unspecific GFP expression in Kenyon cells (see **Sup. Fig. 1d**). Scale bar = 50 µm. **c)** Left: Schematic of the AL highlighting glomeruli labelled by the transgenes shown on the right (same for panels **d,e**). Middle: Comparison of *GMR86C10-Gal4* driven GFP expression in *D. melanogaster* and at two *attP* sites in *D. sechellia*. Arrow: Cell bodies of VM5d (and VM5v) PNs in the anterior-dorsal region above the AL. Note that the relative position of VM5d is more central in *D. sechellia* compared to *D. melanogaster;* its volume is 2.5-fold increased in *D. sechellia*^29^. Sparse labelling of other cells in the brain of both species. Scale bar = 50µm. Insets: AL glomeruli labelled by the respective transgene (same in panels **d,e**). Scale bar = 5 µm. **d)** Labelling of the DM2, DA1, and DL1 glomeruli by a split-Gal4 combination in *D. melanogaster*. In *D. sechellia*, integration of both hemidrivers at the same location (middle) or at different locations (right) leads to very similar expression. 2-3 additional glomeruli (asterisks) are weakly labelled by both transgene combinations. Scale bar = 50 µm. **e)** Split-Gal4 transgenics labelling ventral projection neurons (vPNs) with a more complex dendritic branching pattern in the AL innervating the DM2, DM5 and VM5d glomeruli in *D. melanogaster* (middle) and *D. sechellia* (right). Black arrowhead: unspecific GFP expression in Kenyon cells. Scale bar = 50 µm. **f)** Schematics of the stochastic labelling approach in *D. sechellia*. Top: Timeline of PN development. The first PNs of the adult olfactory system are specified in L3 larvae and additional populations are added by neuroblast divisions during pupal development^62^. Indicated are the time points of heat-shock mediated induction of Flp recombinase expression. L3 = larval stage 3, APF = after pupa formation. Bottom: Flp recombinase mediates stop cassette excision and restores functional reporter expression in few cells (see also **Sup. Fig. 2**). **g)** Left: Anti-FLAG and anti-HA immunofluorescence showing labelling of PNs in a transgenic *D. sechellia* fly carrying two flp-out transgenes, the broad PN driver *VT033006-Gal4* and a Flp recombinase source. MB = mushroom body, LH = lateral horn, AL = antennal lobe. Scale bar = 50 µm. Right: Anti-FLAG immunofluorescence of individual PNs in each brain hemisphere after Flp recombinase mediated cassette excision.

Given the direct link between circuit wiring, signal processing and innate behaviours, we aimed to investigate how the neuroanatomy of individual olfactory PNs, connecting peripheral OSNs to the LH, has evolved lineage-specifically in *D. sechellia*. To this end, we expanded the genetic toolset in *D. sechellia*^29,33,35–37^ to facilitate the comparative analysis of homologous circuits in the central brain at single cell resolution. Our quantitative analysis of PNs between *D. sechellia* and *D. melanogaster* points towards specific rewiring of central brain circuits involved in olfactory behaviours. These results indicate that anatomical rewiring and spatial re-organisation of host cue-related circuits play an important role in olfactory signal processing and complement physiological and morphological alterations in peripheral sensory neurons.

## Results

### Establishment of PN labelling tools in *D. sechellia*

To enable the comparison of homologous PN cell types in the central brain, we first developed several necessary transgenic tools in *D. sechellia*. In *D. melanogaster*, libraries of promoter-Gal4 and split-Gal4 transgenes inserted at defined genomic locations^38,39^ have granted genetic access to numerous cell types with single cell resolution^40–42^. Genetic targeting offers the additional advantage over dye filling and photoactivation experiments^31,34^ that brain images from immunohistochemical staining can be registered on a reference template for quantitative comparison of subtle morphological alterations^43^. However, to achieve genetic labelling with a high degree of precision, site-directed transgenesis at multiple reliable locations in the genome is required to combine several transgenes in the same animal.

We previously found that a broad PN driver from *D. melanogaster* can work well when integrated into the *white* locus of *D. sechellia* (**Fig. 1b**, Ref.^33^) but this site is disadvantageous because it disrupts this gene’s function when using homozygous insertions (resulting in visually impaired flies). To establish other suitable landing sites, we introduced *attP* sequences into the *D. sechellia* genome via random piggyBac-integration and CRISPR/Cas9-mediated homologous recombination at the equivalent locations of well-characterised *attP* sites of *D. melanogaster* and *D. simulans* (*JK22C*^44^, *attP40*^42^, *attP2*^45^, *su(Hw)attP2*^46^, *Dsim1029*^47^) (**Sup. Fig. 1a,b, Sup. Table 1**).

To assess these *D. sechellia* landing sites we attempted integration of a PN Gal4 transgene from *D. melanogaster*, *GMR86C10-Gal4*^48^, for two reasons: first, this driver is sparsely expressed in just two PN populations (innervating the VM5d and VM5v glomeruli^23^) and thereby potentially a more sensitive reporter of the (in)sensitivity of a landing site to genomic positional effects; second, the VM5d PNs receive input from OSNs expressing the receptors Or85c/b, which define a behaviourally important olfactory pathway^29,33^. We succeeded in integrating the *GMR86C10-Gal4* driver into two different *attP* sites (*Dsec-attP40* and *Dsec-attP1029*; **Sup. Table 2**). When these drivers were combined with *UAS-effector* constructs we observed faithful reproduction of the *D. melanogaster* expression pattern, labelling both VM5d and VM5v PNs in the *D. sechellia* brain (**Fig. 1c**). Moreover, the transgenic labelling of VM5d in *D. sechellia* confirmed the expected two- to three-fold increase in size of this glomerulus^29,33,49^ and a change in its relative position within the AL towards the brain midline^49^.

To target PNs more sparsely and to prevent ectopic expression in other brain neurons, we turned to the split-Gal4 system^50^. We selected the *VT033006-BD* (as the *VT033006-Gal4* transgene drives strong and robust expression in PNs, **Fig. 1b**, Ref.^33^) and *GMR72C11-AD* hemidrivers necessary for Gal4 reconstitution. In *D. melanogaster*, this enhancer combination labels PNs innervating the DM2, DA1 and DL1 glomeruli (**Fig. 1d**, SS01867, Ref.^24^). With our new transgenes in *D. sechellia*, we detected reproducible expression in all three homologous glomeruli (**Fig. 1d**, **Sup. Fig. 1c, d**). Integration of one hemidriver into another chromosomal target site (*Dsec-attPpBac-A26*; to avoid potential transvection^51^) (**Fig. 1d**) resulted in a similar expression profile. Last, we tested a second hemidriver combination (SS00587, Ref.^52^) specific for ventral PNs (not targeted by the reagents described above) which led to comparable expression patterns across species (**Fig. 1e**). In sum, we established a series of *attP* sites for site-directed transgenesis in *D. sechellia* and demonstrate that *D. melanogaster* enhancer constructs integrated at these locations can label homologous PNs in *D. sechellia*.

While the split-Gal4 system offers a high degree of cell type specificity, often no hemidriver combination exists for particular PN cell types or multiple cells of the same population are labelled. Therefore, we generated transgenic lines for stochastic cell labelling in *D. sechellia* based on Flp recombinase-mediated excision of a stop cassette (**Fig. 1f**, Ref.^53^). We placed two transgenes through CRISPR-mediated integration via homologous recombination at the location of two *attP* sites, *Dsec-attP40* and *Dsec-attP1029,* on two different chromosomes (II, III, **Sup. Fig. 1b, Sup. Fig. 2a**). As a source of recombinase activity, we generated a transgenic line expressing Flp recombinase (Flp1^54^) under the control of a heat-shock promoter at the *Dsec-attPpBac-A26* site (chromosome IV), which allows combination of all three transgenes without the need of recombination. To test these lines and to investigate the neuroanatomy of PN populations in *D. sechellia*, we sampled PN subtypes by combining our stochastic labelling tools with the broad *VT033006-Gal4* (**Fig. 1b**) and *VT033008-Gal4* transgenes^33^. In transgenic animals carrying either both or a single flp-out cassette, we provided heat-shock-induction of Flp1 after terminal differentiation of PNs at late larval stages and during pupal development (**Fig. 1f**), analogous to previous studies in *D. melanogaster*^18–20^. For both transgene combinations, we generated PN specific labels in female *D. sechellia* and isolated brains where multiple or only individual PNs were labelled (**Fig. 1g**). Together, these new tools provide the opportunity to genetically target and reconstruct single neuron morphologies in *D. sechellia*.

### Single-cell reconstruction of *D. sechellia* PN morphologies

To generate a comprehensive atlas of single cell PN morphologies in *D. sechellia*, we dissected more than 800 brains of Flp-out animals subjected to heat-shock. 620 of these showed strong reporter staining in PNs and the brain neuropil. We selected 184 brains with sufficiently sparse labelling to identify and fully trace individual PNs. We further filtered stained brains by requiring their successful registration to a *D. sechellia* reference brain so that traced cell morphologies could be compared in the same template space. We obtained a dataset of 133 high-quality PNs in female *D. sechellia* from 117 different brains (**Fig. 2a**). We manually traced the morphology of these PNs, and annotated them depending on their dendritic branching pattern following the conventions established in *D. melanogaster*^15^: uniglomerular (uPNs) – comprising uni- and uni^+^-PNs that innervate a unique glomerulus or a unique glomerulus with some additional branches in other glomeruli, respectively – and multiglomerular PNs (mPNs), comprising oligo-, and multi-glomerular types that innervate several or the majority of glomeruli, respectively^15^. In total, our dataset encompasses 93 uPNs and 40 mPNs (**Fig. 2a, Sup. Fig. 2b**).

**Figure 2.**
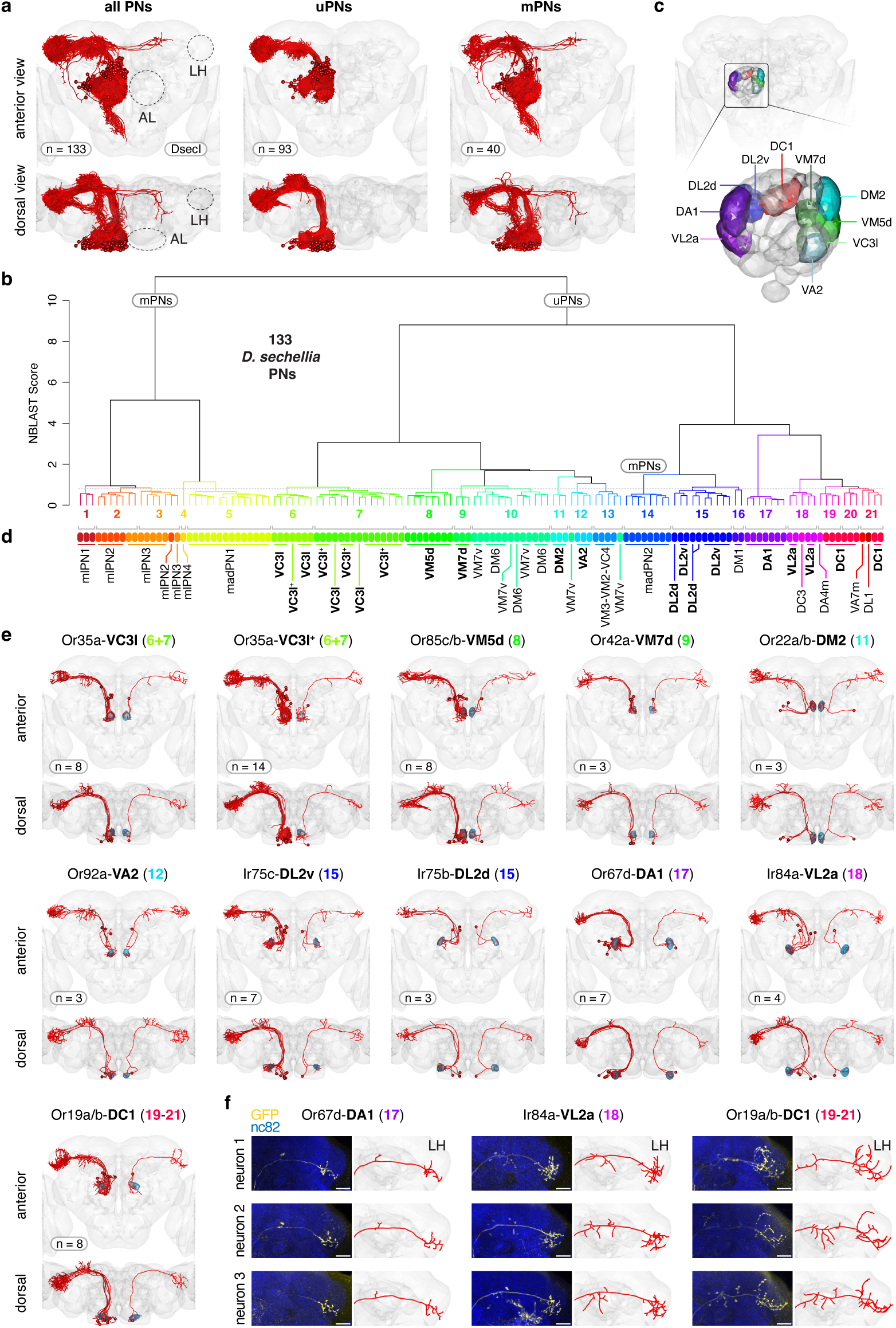
Identification of PN cell types in *D. sechellia*. **a)** Right: 133 traced PNs registered on the *D. sechellia* DsecI reference brain. Middle: 93 uniglomerular and uni^+^-glomerular PNs (uPNs). Left: 40 multiglomerular PNs (mPNs). Top: anterior view, bottom: dorsal view. A full list of all traced PNs is provided in **Sup. Table 3**. **b)** NBLAST clustering of the *D. sechellia* PN dataset results in 21 distinct clusters. **c)** *D. sechellia* reference brain with segmentation and annotation of AL glomeruli shown in panel **e**. **d)** Manual curation of the 21 NBLAST clusters identifies 26 PN types. Colours indicate PN type, labels indicate the (main) innervated AL glomerulus. In bold: uPN types shown in panel **e**. For VC3l, PNs with restricted innervation to VC3l or with additional innervations of other glomeruli (VC3l^+^) are shown. **e)** Eleven *D. sechellia* uPN types. Labels indicate the olfactory receptor expressed in OSN partner neurons, the main innervated glomerulus (bold) and the cluster number of the NBLAST result (as in panel **b**). Left brain hemisphere: All neurons of the respective type in the *D. sechellia* dataset; right brain hemisphere: individual, representative neuron. Blue: segmentation of the AL glomerulus/i with main dendritic innervation. The sample size for each type is indicated. **f)** Anti-HA immunofluorescence of three representative Or67d-DA1, Ir84a-VL2a, and Or19a/b-DC1 PN axons (left) and the corresponding tracings (right) with stereotypic branching in the LH. Scale bar = 20 µm.

To provide additional categorisation of *D. sechellia*’s PNs, we used NBLAST, which compares the morphological properties of traces and clusters them hierarchically^55^. Twenty-one distinct cell types emerged from this analysis (**Fig. 2b**). To further improve this clustering-based characterisation, we segmented AL glomeruli in the *D. sechellia* reference brain (based on Refs.^34,56^) (**Fig. 2c**) and used the dendritic innervation pattern of PNs to assign them to distinct AL glomeruli. In combination with the NBLAST output, this led to a refinement of our annotation, resulting in 26 distinct PN cell types in our *D. sechellia* PN dataset (**Fig. 2d**, **Sup. Fig. 2c**).

Earlier studies in *D. melanogaster* have shown that PNs from individual glomeruli project to specific sub-regions of the LH and show a stereotypic branching pattern across animals^18–20^. This morphology suggests that connectivity with downstream LH neurons and odour representation in the LH is conserved among individuals (and genetically hardwired, in contrast to PN-Kenyon cell connections in the MB that rather follow a stochastic connectivity pattern)^34^. Comparing individual uPN neurons of the same type, we confirmed this stereotypy exists also for *D. sechellia*’s PNs (**Fig. 2e, f**) indicating that the genetic hardwiring of cell morphology (and potential synaptic connectivity) among PNs is a conserved feature across these drosophilids.

The representation of PNs in our dataset was not uniform: for several PN types, we obtained only single representatives while for others we recovered up to fourteen examples (**Sup. Fig. 2c**). This bias might be caused by the genetic driver we used, which labels many but not all antero-dorsal and lateral PN populations. Moreover, the time window chosen for heat-shocks (late larval and early pupal development, **Fig. 1f**) might favour recombination in selected PN neuroblasts depending on their birth order^24^. Despite these caveats, the dataset contains a broad diversity of PN types with sufficient coverage to enable to compare single representatives of the same cell type intra-specifically, and between species.

### *D. sechellia* has evolved different PN morphology in noni-sensing pathways

To compare *D. sechellia* PNs to homologous neurons in *D. melanogaster*, we first transferred PN reconstructions from *D. melanogaster* light imaging data (the Virtual Fly Brain^57^) and two electron microscopy connectomes^15,16^ into the *D. melanogaster* IS2 reference space (**Fig. 3a**). By limiting our analysis to cell types of the anterior-dorsal and lateral PN clusters with at least three representatives within our *D. sechellia* dataset (19/26 cell types), we were able to systematically compare homologous cell types at single cell resolution between both species.

**Figure 3.**
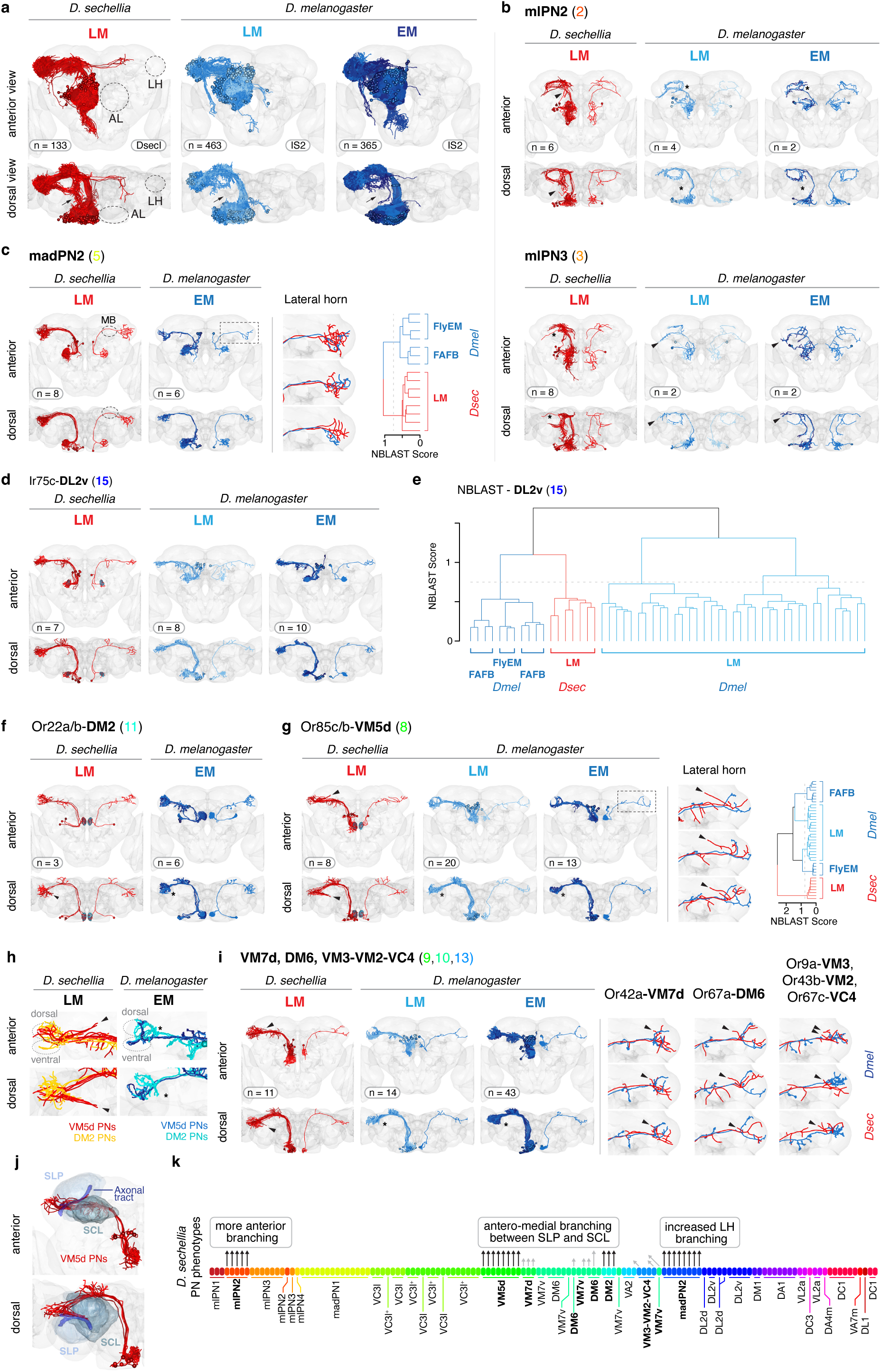
Comparison of *D. sechellia* and *D. melanogaster* PN morphologies. **a)** Left: *D. sechellia* light microscopy (LM) dataset of 133 PNs (with soma in the anterior-dorsal or lateral PN cluster). Middle: *D. melanogaster* LM dataset (from the Virtual Fly Brain^57^) of 463 ad/l PNs. Of these, 285 neurons were annotated, 178 lacked annotations. Right: *D. melanogaster* electron microscopy (EM) data^15,16^ of 365 ad/l PNs (317 uPNs, 48 multi PNs). Note that in the *D. melanogaster* datasets the soma position is mis-placed in some cases. AL = antennal lobe, LH = lateral horn. Top: anterior view, bottom: dorsal view. Black arrow: increased anterior branching in mPNs of *D. sechellia* (see panel **b**). **b)** Two types of mPNs with cell soma in the lateral cell cluster (mlPNs) in *D. sechellia* (left) and *D. melanogaster* (middle and right). mlPN2 display axonal branches reaching towards the anterior brain (arrows) only in *D. sechellia*; mlPN3s in *D. sechellia* show less branching into the LH (asterisk). Here and in the following panels, all neurons of a certain type are shown in the left hemisphere; a single, representative example neuron in the right hemisphere. **c)** mPNs with anterior-dorsal cell body position (madPN1) almost completely lack mushroom body (MB) innervations and show more complex axonal arbours in the *D. sechellia* LH (right: overlay in the *Dmel* IS2 reference space). NBLAST clustering separates *D. sechellia* (*Dsec*) PNs from their *D. melanogaster* (*Dmel*) homologs. **d)** Dl2v uPNs show a conserved morphology between both species (see **Sup. Fig. 3** for other PN types). **e)** NBLAST does not separate *D. sechellia* Dl2v PNs from their *D. melanogaster* homologs supporting the conserved neuroanatomy of this cell type. **f)** Comparison of DM2 uPN morphologies between *D. sechellia* (left, LM) and *D. melanogaster* (right, EM) confirms published observations of additional antero-medial PN branching in *D. sechellia*^29^ (arrowhead, absent in *D. melanogaster*: asterisk). **g)** VM5d PNs in *D. sechellia* (left) display a prominent antero-medial branch (arrowhead) absent in *D. melanogaster* (asterisk). Right: Three representative examples of LH branching of VM5d PNs in both species (overlay in the *Dmel* IS2 reference space). NBLAST clustering separates *D. sechellia* (*Dsec*) PNs from their *D. melanogaster* (*Dmel*) homologs. **h)** Visualisation of VM5d and DM2 PNs in the IS2 reference space for *D. sechellia* (left) and *D. melanogaster* (right). Indicated are the dorsal and ventral LH areas (circles). Arrowhead = antero-medial branching of both populations in *D. sechellia* which is absent in *D. melanogaster* (asterisk). **i)** A few other uPNs show branching into the DM2/VM5d innervated antero-medial brain area in *D. sechellia* (arrowhead) absent in *D. melanogaster* (asterisk). Right: Individual examples of VM7d, DM6 and VM3-VM2-VC4 uPN branching in the LH (overlay in the *Dmel* IS2 reference space). **j)** Anatomical regions of the dorsal *D. melanogaster* brain. The *D. sechellia* specific branch of VM5d PNs elongates along an axonal tract between the superior lateral protocerebrum (SLP) and the superior clamp (SCL). **k)** Summary of the observed PN morphological changes in *D. sechellia* compared to *D. melanogaster*. Arrows indicate if individual neurons display the phenotype described on top. Arrow length gives an indication of the strength of the observed phenotype (longer arrow = stronger phenotype; black arrows = 100% penetrance of phenotype).

We first focused on mPNs, which form a heterogenous group with complex cell morphologies (**Fig. 3b, c**). We identified five different mPN types in *D. sechellia*. Of these, three are housed in the lateral PN cell cluster (“multi-lateral PN 1-3”, mlPN1-3) and two display alterations in their branching phenotypes in *D. sechellia* compared to *D. melanogaster* (mlPN2, mlPN3, **Fig. 3b**): mIPN2s send more branches into anterior brain areas in *D. sechellia,* mIPN3s instead show a reduced branching phenotype in the LH in this species with more pronounced projections towards the brain midline (**Fig. 3b**). mlPN1’s morphology is comparable across species (**Sup. Fig. 3a**).

The two cell types with antero-dorsal soma position, madPN1 and madPN2 (“multi antero-dorsal PN”), show a similar overall branching pattern (**Fig. 3c, Sup. Fig. 3b**). madPN2 neurons, however, which innervate the DL2d and DL2v glomeruli, form a more complex axonal arbour in the *D. sechellia* LH (**Fig. 3c**). These neurons bypass the MB calyx without visible bouton formation (one branch was detected in one *D. melanogaster* neuron), indicative of only a minor role (if any) in associative learning. These neurons receive input from acid-sensing Ir75c and Ir75b OSNs, of which the latter play a role in species-specific oviposition site preference^58^. The altered branching of this population in the LH could lead to differential weighting of acid-related cues in *D. sechellia* and have an impact on egg-laying behaviour. To confirm the visually observed difference between *D. sechellia* and *D. melanogaster* neurons, we compared neuronal morphologies using inter-specific NBLAST after transfer of all PN traces into the same IS2 reference space through bridging registrations. *D. melanogaster* madPN2s formed a distinct cluster from those in *D. sechellia*. This clustering likely reflects biological rather than technical differences across datasets because, for other cell types, light and electron microscopy samples co-clustered (**Fig. 3g**) and highly similar cell types did not segregate by either species or technique (**Fig. 3d,e**). While we detected some morphological variation in mPNs, it is currently challenging to link the observed modifications to function because mPNs pool input from many OSN populations.

We next focused on the morphology of (excitatory) uPNs that were unambiguously identified by their dendritic AL innervation pattern and main OSN partners^15^. In eight uPN types, we did not detect any obvious morphological difference between both species, supporting the overall conserved anatomy of many PN types in *D. sechellia* (**Fig. 3d,e, Sup. Fig. 3c**). Despite the broad conservation, however, we did detect several inter-specific differences. Consistent with published work^29^ *D. sechellia* DM2 PNs form an axonal branch, which is absent in *D. melanogaster*, leaving the densely packed LH area towards the antero-medial area of the brain (**Fig. 3f**). The same antero-medial brain area is also innervated by multi-glomerular PNs in both species which form more projections in the anterior part of the brain in *D. sechellia* than in *D. melanogaster* (**Fig. 3a**).

Notably, VM5d PNs display an even more prominent antero-medial branch, innervating the same brain region as DM2 PNs in *D. sechellia* (**Fig. 3g,h**). Remarkably, while within DM2 the overall axonal morphology within the LH is conserved, VM5d PNs display a clear reduction in their LH branching in *D. sechellia*. They maintain their innervation of the dorsal LH zone, restricted to olfactory inputs, while their branches in the ventral area, which receives multimodal input^15,16,59^, is reduced (**Fig. 3g**). The co-wiring of both PN populations out of the LH is particularly interesting as the cognate presynaptic sensory neuron partners of DM2 and VM5d PNs (Or22a and Or85c/b OSNs) are both involved in long-range attraction to the host fruit of *D. sechellia*^29,33^. The new antero-medial branch in both PN populations could therefore represent an additional evolutionary novelty in *D. sechellia* setting these two pathways further apart from other PN populations. To investigate this observation further, we performed photoactivation experiments in the antero-medial branch area in *D. sechellia* in an effort to label all AL glomeruli innervated by PNs that contribute to this structure. We found that only DM2 and VM5d PNs, which are both located in the dorso-medial AL, were reverse labelled by this approach, further indicating that these two populations are the main contributors to the *D. sechellia*-specific innervations of this brain area (**Sup. Fig. 4a**). Upon further inspection, we detected a few other PN types in *D. sechellia* (DM6, VM7v, VM7d and VM2) which also display axonal processes in this same brain area with low frequency (3/8 in DM6, 2/6 in VM7v) or with a small antero-medial branch (VM7d, VM2) (**Fig. 3i**). All these PN populations respond to food-related odours^56^ and their respective glomeruli are located in close proximity to DM2 and VM5d in the AL (**Fig. 4c**). However, of all eleven uPN types sampled in our dataset, only DM2 and VM5d PNs consistently displayed novel projections into the antero-medial brain area.

**Figure 4.**
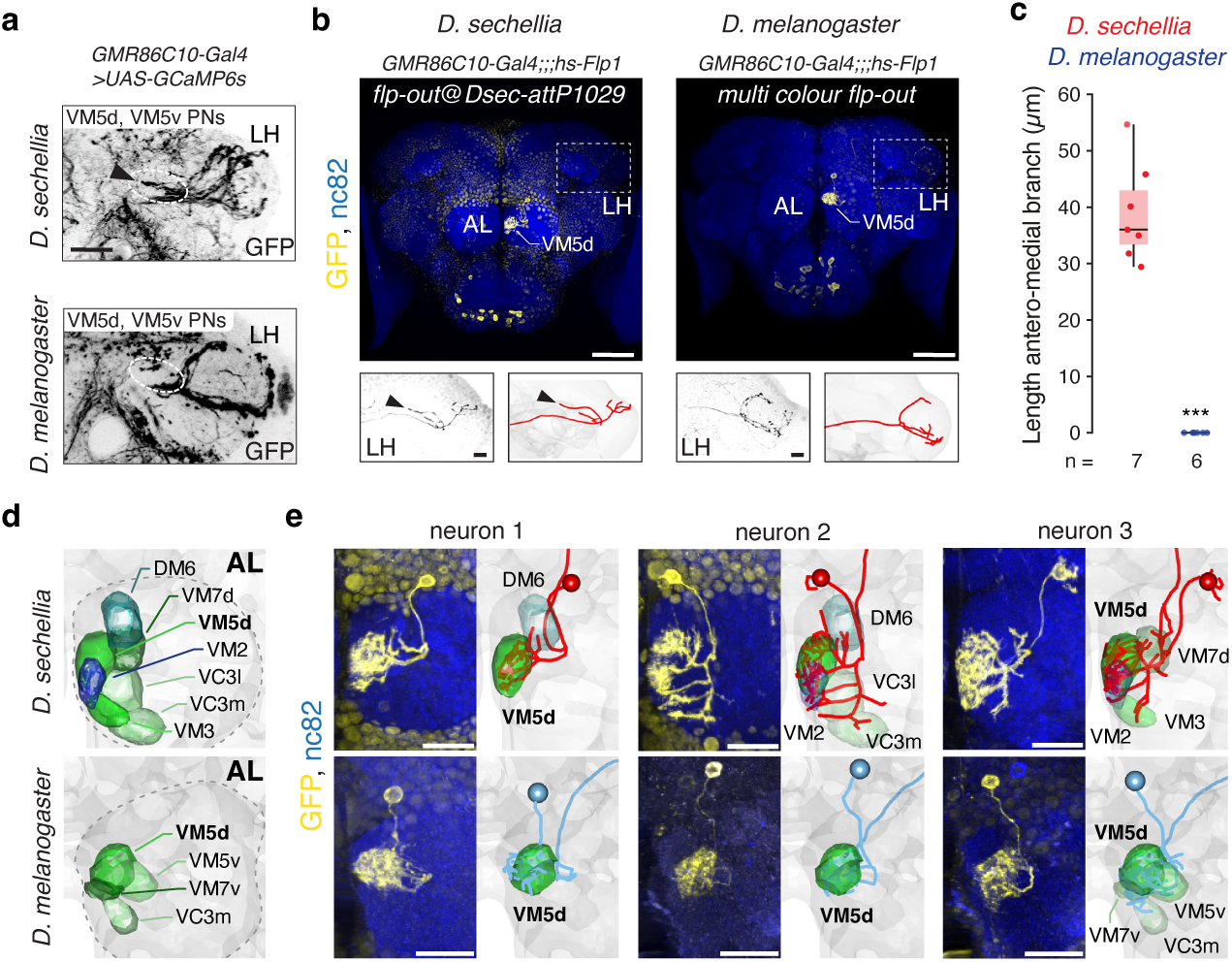
Novel axonal and dendritic branching in VM5d PNs. **a)** Anti-GFP immunofluorescence showing the arborisation of PNs in the *GMR86C10-Gal4* transgenic line (Fig. 1c) in *D. sechellia* and *D. melanogaster.* Circle and black arrowhead: antero-medial branch of VM5d PNs in *D. sechellia*. LH = lateral horn. Scale bar = 20 µm. **b)** Anti-FLAG immunofluorescence in transgenic *D. sechellia* (left) and *D. melanogaster* (right) flies carrying flp-out reagents and the *GMR86C10-Gal4* transgene. Nc82 is a broad neuropil marker. Scale bar = 50µm. Below: axonal branching of one VM5d PN and neuron tracing. Arrowhead: antero-medial branch of VM5d PNs in *D. sechellia.* Scale bar = 10 µm. **c)** Quantification of VM5d PN antero-medial branch length between species (n = 6-7 females) (pairwise Wilcoxon rank-sum test, two-sided): *** *P* < 0.001. **d)** AL segmentation of selected glomeruli neighbouring VM5d in *D. sechellia* (top) and *D. melanogaster* (bottom). AL = antennal lobe. **e)** Left: Anti-FLAG immunofluorescence of individual VM5d PNs in the *D. sechellia* (top) or *D. melanogaster* (bottom) AL. Right: reconstruction of the dendritic arbour of the stained neuron shown on the left and VM5d and other innervated glomerular boundaries. Scale bar = 20 µm. Three neurons are shown for each species.

To further characterise the target area of DM2 and VM5d PNs, we transferred anatomical segmentations from the Virtual Fly Brain to the IS2 reference space (**Fig. 3j**). The *D. sechellia* specific antero-medial PN branches elongate along an axonal tract between the superior clamp and the superior lateral protocerebrum. This tract is innervated by mPNs, mushroom body output neurons, lateral horn neurons and others in *D. melanogaster* (see below) and represents a pre-defined exit point of the LH.

Together, the co-registration of our *D. sechellia* PN dataset with existing traced neurons in *D. melanogaster* and our high-resolution analysis have revealed several morphological differences in mPN and uPNs. In support of a role in modified olfactory processing, we detected several instances of circuit rewiring in *D. sechellia* either in the LH (madPN2, VM5d) or towards a novel target area in the antero-medial brain innervated by olfactory pathways detecting noni odours (DM2 and VM5d) (**Fig. 3k**).

### *D. sechellia* VM5d PNs display differences in both axonal and dendritic wiring

To confirm the novel antero-medial branching of VM5d PNs in *D. sechellia* with another genetic driver line, we made use of the *GMR86C10-Gal4* transgene (**Fig. 1c**). This transgene allows the reproducible labelling of all VM5d PNs (eliminating any potential bias in our stochastically-labelled neuron dataset). Using this independent driver, we again detected the novel antero-medial branching of VM5d PNs (**Fig. 4a**). By combining this driver with our stochastic labelling tools, and using equivalent transgenes in *D. melanogaster*, we reconstructed additional VM5d PN morphologies in both species. All of these (n=7) display a prominent antero-medial branch leaving the LH neuropil in *D. sechellia* which is absent in *D. melanogaster* (**Fig. 4b, c**), further supporting the penetrance of this phenotype in *D. sechellia*.

Within the AL, the VM5d glomerulus is more medially located and two- to three-fold larger in *D. sechellia* compared to *D. melanogaster* (or *D. simulans*) (**Fig. 4d**)^29,33,34,49,56^. Upon close inspection of VM5d dendritic branches in the AL of both species, we detected several instances of unexpected dendritic targeting in *D. sechellia*. VM5d PNs frequently (in 4/7 examples) did not innervate a singular glomerular target but dendrites also branched into the DM6, VM7d/v, VM2 or VC3 glomeruli (**Fig. 4e**). In *D. melanogaster*, such targeting promiscuity was not detected, with VM5d dendritic branches showing minor innervation of VM7d or VC3m only in few cases (2/6). Intriguingly, the glomeruli innervated by *D. sechellia* VM5d PNs are the same those whose PNs show occasional axonal antero-medial branching and co-wiring with VM5d PNs. In sum, this PN type does not only show axonal but also dendritic wiring alterations in *D. sechellia*.

### Antero-medial PN axon targeting is a *D. sechellia*-specific, recessive trait

We next wanted to clarify if the novel antero-medial branches of DM2 and VM5d PNs are *D. sechellia* specific and therefore turned to its closest relative *D. simulans.* To visualise PNs in this species, we first generated a *D. simulans VT033006-Gal4* line and combined it with a fluorescent reporter^60^. We imaged the LH and adjacent brain regions and compared PN innervation patterns across all three species. Only *D. sechellia* shows staining in the area antero-medial to the LH (**Fig. 5a**). Next, we expressed photoactivatable GFP^34^ in *D. simulans* pan-neuronally. Following specific activation of the VM5d glomerulus in *D. simulans*, we detected no branches reaching into the antero-medial brain area next to the superior clamp as seen in *D. sechellia* (**Sup. Fig. 4b**).

**Figure 5.**
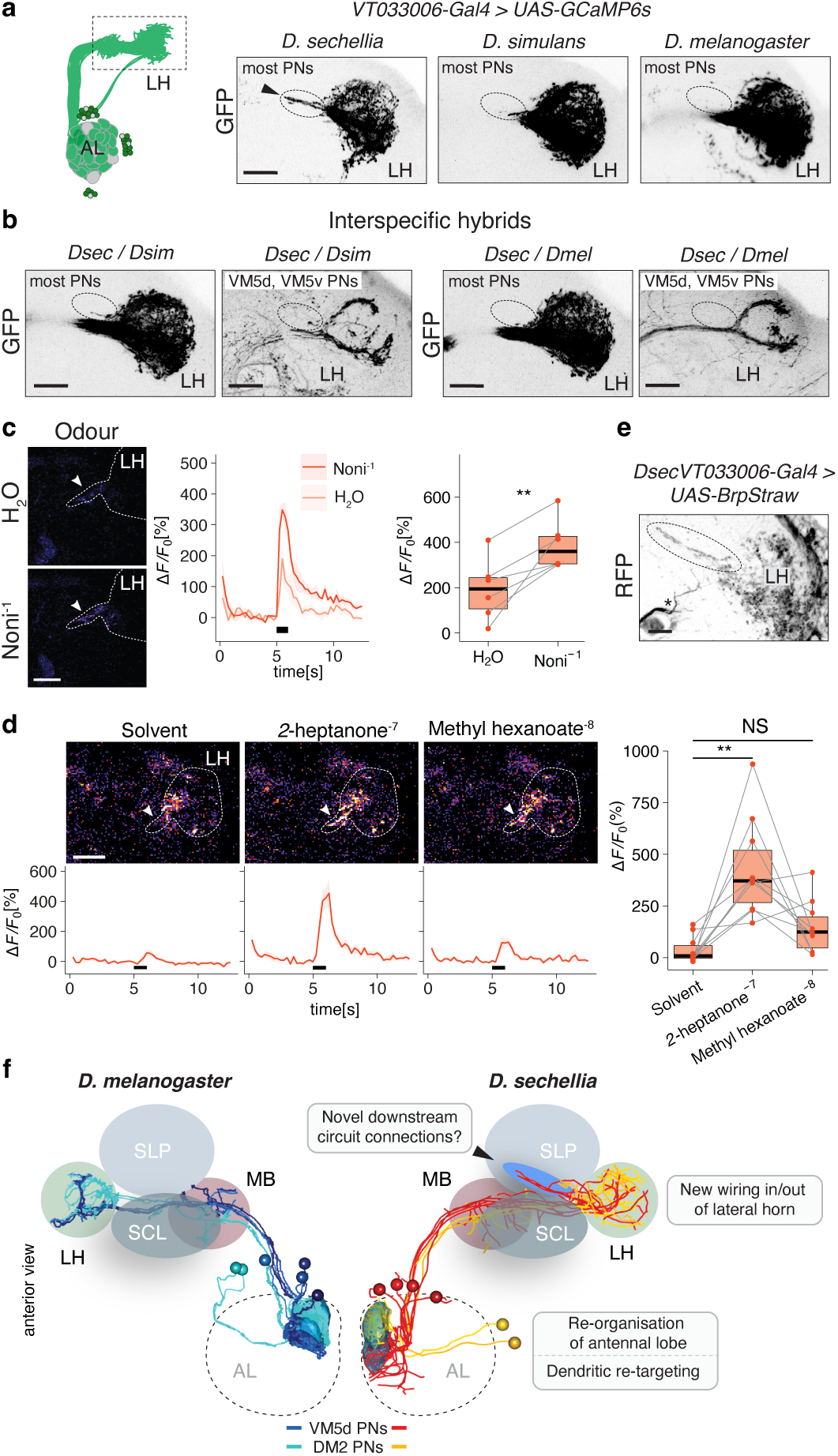
Genetic basis and activity of *D. sechellia*-specific PN branches. **a)** Left: schematic of labelled PNs. Right: Anti-GFP immunofluorescence in *VT033006-Gal4, UAS-GCaMP6s* transgenics of three species. Only in *D. sechellia*, axonal branching from the LH towards the antero-medial part of the brain is detected (brain midline to the left). **b)** Anti-GFP immunofluorescence showing PN branching of transgenic interspecific hybrids expressing *VT033006-Gal4, UAS-GCaMP6s* (left) or *GMR86C10-Gal4*, *UAS-GFP* (right). Left: *D. sechellia*/*D. simulans* hybrids. Right: *D. sechellia*/*D. melanogaster* hybrids. No *D. sechellia* equivalent antero-medial branch in VM5d or other labelled PNs is detected (circle). Scale bar = 20 µm. **c)** Left: Odour-evoked neuronal activity in the *D. sechellia* antero-medial PN branch in *VT033006-Gal4, UAS-GCaMP6f* transgenic animals upon presentation of water and noni juice (at 10^-1^ dilution). White arrow: branch area quantified. Scale bar = 25 µm. Middle: Relative increase in fluorescence (ΔF/F) in percentage upon stimulus presentation. Black bar = stimulation window. Right: Maximal responses towards each stimulus (n = 6 females). For these and box plots in panel **d**, the centre line represent the median, the box bounds represent the first and third quartiles, and whiskers depict at maximum 1.5× the interquartile range; individual data points are overlaid. Paired t-test, two-sided: \*\**P* < 0.01. **d)** Odour-evoked neuronal activity in the *D. sechellia* antero-medial PN branch in *VT033006-Gal4, UAS-GCaMP6f* transgenic animals upon presentation of solvent (= dichloromethane), heptanone^-7^ or methyl hexanoate^-8^. The latter two specifically activate VM5d and DM2 PNs at the given concentration^33^. White arrow: branch area quantified. Scale bar = 25 µm. Below: Relative increase in fluorescence (ΔF/F) in percentage upon stimulus presentation. Right: Maximal responses towards each stimulus (n = 10 females). Paired t-test, two-sided: \*\**P* < 0.01, NS = *P* > 0.05. **e)** Expression of a *Brp^Straw^*transgene in *D. sechellia* PNs reveals labelling of potential pre-synaptic sites in the antero-medial branch area (in circle). Asterisk = unspecific labelling. Scale bar = 10 µm. **f)** Summary of anatomical differences in the AL – LH circuitry between *D. melanogaster* and *D. sechellia*. Shown are four VM5d and two DM2 PNs in both species. *D. sechellia* shows a re-organisation of its AL glomeruli positions and dendritic re-targeting in VM5d PNs. In the LH, less axonal branches are present in the ventral part with antero-medial branches (arrowhead) exiting the LH (and co-wiring of VM5d and DM2 PNs), potentially forming novel connections with downstream partners. MB = mushroom body, SLP = superior lateral protocerebrum, SCL = superior clamp, LH = lateral horn, AL = antennal lobe.

To gain insights into the genetic architecture of the *D. sechellia* specific antero-medial PN branching, we analysed PNs in *D. sechellia*/*D. simulans* and *D. sechellia/D. melanogaster* hybrids. In both hybrid combinations, no *D. sechellia*-like branching phenotype is detected (with a minimal branch observed in the *D. sechellia/D. simulans* hybrids in few (2/6) cases) (**Fig. 5b**). Consistent with this result, in inter-specific hybrids carrying VM5d/VM5v specific genetic labels using the *GMR86C10-Gal4* driver, no antero-medial branch was present (**Fig. 5b**). Collectively, these experiments suggest that altered PN axonal branching into the antero-medial brain is a derived, recessive trait in *D. sechellia*.

### Noni odour-evoked activity is transmitted to a new target area

To test if the *D. sechellia* DM2 and VM5d antero-medial PN branches transmit noni odour-evoked activity, we presented olfactory stimuli to *D. sechellia* flies and performed 2-photon calcium imaging using the broad PN driver *VT033006-Gal4* coupled with *UAS-GCaMP6f*. Compared to control stimulation with water, we observed a robust noni response in *D. sechellia* in the antero-medial branch area (**Fig. 5c**). This result supports that food-related olfactory signals converge into this target area and are potentially transmitted to downstream partners. To monitor the contribution of both individual PN cell populations, VM5d and DM2, to the observed activity pattern, we chose *2*-heptanone and methyl hexanoate at low concentrations as stimulants which mainly activate Or85c/b (VM5d) and Or22a/b (DM2) sensory neurons in the antenna, respectively^33^. For both stimuli, we could detect activity in the *D. sechellia*-specific branching area (**Fig. 5d**), however to a lesser degree for the Or22a/b specific odour, consistent with the less prominent innervation of these neurons compared to VM5d PNs.

Finally, we labelled synaptic active sites in these neurons using a *UAS-Brp-short^mStraw^* transgene^61^. Like axons within the LH, the antero-medial PN branches show distributed punctate staining throughout their whole length (**Fig. 5e**). This result suggests that pre-synaptic sites are established within these branches, consistent with them forming functional synapses with downstream neurons (see Discussion). Together with the above physiological results, these experiments support our anatomical data and indicate that noni-responsive pathways form *D. sechellia*-specific connections in the antero-medial brain and suggest that they have functional relevance for the interpretation of noni odours in downstream circuits.

## Discussion

We have used two closely-related, genetically-accessible fly species as a powerful paradigm to understand principles of neural circuit evolution. Using a suite of novel genetic tools, we were able to genetically compare more than twenty homologous cell types in the central brain of multiple individuals. Olfactory PNs are ideally suited for this approach as they are critical for olfactory processing and can be unambiguously identified through the stereotypic wiring of the olfactory system.

The *attB/attP* system that we have employed offers the advantage that published constructs of *D. melanogaster* can be re-used to label homologous cell types across the *D. melanogaster/D. simulans/D. sechellia* trio. The four *attP* sites described here, located on each of the four chromosomes, allow the combination of up to four transgenes without the need of recombination. Future efforts will be necessary to transfer other markers into *D. sechellia* to facilitate the creation of more complex genotypes.

Our analysis of PN morphologies across *D. melanogaster* and *D. sechellia* revealed that while most cell types exhibit a conserved morphology, modifications in multi- and uniglomerular PNs exist. In particular, the axonal rewiring of the DM2 and VM5d PN populations in *D. sechellia* is intriguing as both pathways are involved in long-range attractive behaviour towards noni in *D. sechellia*^29,33^. The neuron types are developmentally not closely-related – deriving from distinct neural stem cell lineages^62^ – and the specificity of the antero-medial branch for those two cell types implies that anatomical convergence is an adaptive trait selected by their common function in noni detection. While there is no specific evidence for increased axonal-axonal synaptic contacts between both populations in *D. melanogaster*^15^, these might be present in *D. sechellia* as axon-axon proximity is strongly related to the number of synapse between cells^63^. Querying the *D. melanogaster* connectome, beyond mPNs, many other neurons innervate the antero-medial branch area, including those of the superior lateral, medial and intermediate protocerebrum, mushroom body output neurons, and lateral horn neurons (**Sup. Fig. 5, Sup. Table 4**). The latter represent excellent candidates for novel connections in *D. sechellia* as they show stereotypic odour responses across animals in *D. melanogaster*^27,28^ and receive on average excitatory input from 6-7 PNs^15,16,28^. In our approach, we sampled PNs across many brains, and it is unclear how many PNs innervate the *D. sechellia* antero-medial branch area per animal. However, the number of DM2 and VM5d PNs per hemisphere (2 and 4, respectively) together with some innervations of other PN types (DM6, VM7d, etc.) nicely matches the average input per LH neuron. These serve as better odour categorisers – in this case for noni as highly ecologically relevant odour mixture – than individual PNs and stereotypically pool related inputs. This coding strategy improves signal-to-noise ratio of LHN responses and could lead to increased sensitivity for noni signals at this processing stage in *D. sechellia*. Our calcium imaging data support that these noni signals are transmitted into the antero-medial branch area as does the presence of pre-synaptic sites. Overall, our data favour a model where convergence of behaviourally relevant olfactory input onto LHNs through rewiring of PN axons leads to species-specific representation of noni in the *D. sechellia* brain (**Fig. 5f**).

Previous work in *D. melanogaster* PN development has shown, that axonal and dendritic wiring is controlled by common *cis*-regulatory networks^64^. In the case of VM5d, where in *D. sechellia* axonal branching in the central brain and dendritic targeting in the AL are modified, both could have a common genetic basis. Recently developed methods to study PN wiring^21–24^ will be instrumental to investigate how species-specific wiring can be established during development. The re-location of VM5d in the *D. sechellia* AL and dendritic innervation of neighbouring glomeruli could allow pooling of food-related inputs into a single pathway. So far, response properties in these PNs have been sampled only with a single odorant^33^. It will be interesting to test, if targeting promiscuity in *D. sechellia* impacts physiological VM5d PN response properties and leads to a broadened response profile in these neurons.

Overall, the genetic toolset presented here opens the door for more sophisticated genetic manipulations in drosophilids. Comparative studies in homologous mating circuits underlying species-specific behaviours^60,65–70^ can build on this approach to delineate neuroanatomical species differences deep insight the brain. In combination with connectomic reconstructions after volumetric electron microscopy imaging^15,16^, it represents a powerful way to identify novel circuit motifs and their plasticity across individuals. In *D. sechellia* other phenotypic traits like modified gustation^71^, oviposition^58^, metabolism^72–74^, immune-response^75^, sleep^76^, circadian rhythms^77^, song production^68^ or walking^78^ will benefit from the reagents presented here. Sampling closely related species points towards the accumulation of several changes in few pathways, from the periphery to the central brain, and studies in other selected drosophilids with diverse phenotypes and evolutionary relatedness will help to detect recurrent evolutionary novelties. This gained knowledge is not only relevant for flies but will eventually inform how brains adapt to changing environments.

## Supporting information

Supplementary Data (Dsec PN traces)

## Author Contributions

T.O.A. conceived the project. All authors contributed to experimental design, analysis and interpretation of results. Specific experimental contributions were as follows: B.R.D performed immunostainings, tracings, and analysed all PN morphological data. E.B. mapped transgenic insertion sites and performed molecular cloning. S.T. performed photo-activation and calcium imaging experiments. J.P. performed immunostainings, helped with molecular cloning and maintained transgenic fly strains. L.A. performed molecular cloning and helped with line establishment. G.L. helped with micro-injections and maintained transgenic fly strains. R.B. provided guidance throughout the project. T.O.A. performed molecular cloning, immunostainings and generated transgenic lines. T.O.A. wrote the paper with input from all authors. T.O.A. and B.R.D. prepared figures. All authors approved the final version of the manuscript.

## Acknowledgements

We thank David Stern, David O’Brochta, Iris Salecker, Stephan Sigrist, Benjamin Prud’homme, Sophie Caron, Gerald Rubin, Yoshi Aso and Heather Dionne for sharing reagents; Yoshi Aso and Greg Jefferis for pointing us to the enhancer-Gal4 and split-Gal4 combinations used in this study and Alexander Bates, Philipp Schlegel and Sebastian Cachero for help with the natverse ecosystem. We also thank Ana Depetris-Chauvin and Silke Sachse for sharing their *D. sechellia* antennal lobe segmentation prior to publication, Anabela Rebelo Pimentel for help with embryo alignments for microinjections, Roman Arguello, Gáspár Jékely and Anne von Philipsborn for comments on the manuscript, and Gáspár Jékely for support to B.R.D.. This work was supported by a Deutsche Forschungsgemeinschaft Walter-Benjamin Fellowship (E.B.), a Marie Skłodowska-Curie Actions Individual Fellowship (836783), an EMBO Long-Term Fellowship (ALTF 454-2019) and a Japanese Society for the Promotion of Science (JSPS) Overseas Research Fellowship (202360258) (S.T.), the University of Lausanne, the Swiss National Science Foundation (SNSF) (310030B-185377), European Research Council Consolidator (615094) and Advanced (833548) grants (R.B.), a SNSF Ambizione (PZ00P3 185743) and Starting Grant (TMSGI3_211391/1), the Fondation Pierre Mercier pour la science, the Novartis Foundation for biomedical research, and the University of Fribourg (T.O.A.).

## Methods

### Data reporting

Preliminary experiments were used to assess variance and determine adequate sample sizes in advance of acquisition of the reported data.

### Drosophila husbandry

*Drosophila* stocks were maintained on standard wheat flour–yeast–fruit-juice medium under a 12-h light:12-h dark cycle at 25 °C. For all *D. sechellia* strains, formula 4-24 instant Drosophila medium, blue (Carolina Biological Supply) soaked in noni juice (Raab Vitalfood) was added on top of the standard food. A detailed list of wildtype and transgenic lines used and generated in this study can be found in **Sup. Table 5.**

### *Drosophila* microinjections

Transgenesis of *D. sechellia* was performed in-house as described^29^ with exception of P-element transgenesis performed by WellGenetics. In details, for *piggyBac* and P-element transgenesis, a *piggyBac* or *P-element* vector (300 ng µl^-1^) and *piggyBac*^79^ or *P-element* helper plasmid^47^ (300 ng µl^-1^) were co-injected. For CRISPR–Cas9-mediated homologous recombination, a mix of a sgRNA-encoding construct (150 ng μl−1) and donor vector (400 ng μl−1) was injected into *Dsec nanos-Cas9*^29^. Site-directed integration into *attP* sites was achieved by co-injection of an attB-containing vector (400 ng μl−1) and pBS130 (400 ng μl−1) (encoding PhiC31 integrase under control of a heat shock promoter (Addgene #26290, Ref.^80^)). All concentrations are given as final values in the injection mix. Insertion efficiencies for *attP* sites are listed in **Sup. Table 2**.

### Molecular cloning

#### piggyBac-attP vector

We generated an *attP* site containing piggyBac vector marked with 3xP3-DsRed expression by replacing the *3xP3-YFP* cassette in *pBac(attP-3P3-YFP)*^47^ (Addgene #86860) with a floxed *3xP3-DsRed* cassette from *pHD-DsRed-attP*^81^ via Gibson Assembly.

#### Donor vectors for homologous recombination

Homology arms (0.8-1.6 kb) for individual target loci were amplified from *D. sechellia* (*Drosophila* Species Stock Center [DSSC] 14021-0248.07) genomic DNA and inserted into the respective donor vector backbone using either restriction cloning or Gibson Assembly. For *Dsec*-*attP2, Dsec-attPJK22C,* and *Dsec-attP40,* homology arms were inserted into *pHD-DsRed-attP*^81^ via restriction cloning.

#### Dsec-su(Hw)attP2

We introduced an FRT site into *pHD-DsRed-attP* by adding its sequence to a PCR amplicon together with NcoI, BamHI overhangs and inserted the digested amplicon into the NcoI/BamHI digested *pHD-DsRed-attP* backbone resulting in *pHD-FRT-DsRed-attP*. Homology arms for the *su(Hw)attP2* locus were amplified from *D. sechellia* (DSSC 14021-0248.07) genomic DNA and inserted into *pHD-FRT-DsRed-attP* via restriction cloning.

#### Dsec-attP1029

We generated a *pHD-mhc-FRT-mFRT71-YFP* donor vector by amplification of three individual PCR amplicons from *pHD-DsRed-attP, pUC57-MHC-DsRed*^82^ and *p3xP3-YFP-attP*^47^. PCR primers included overhangs generating the *FRT-mFRT71* sites and all three amplicons were combined to a single vector via Gibson assembly. Homology arms for the *Dsim1029* homologous locus^47^ were amplified from *D. sechellia* (DSSC 14021-0248.07) genomic DNA and inserted via restriction cloning and Gibson Assembly resulting in *Dsec1029-mhc-YFP*.

#### UAS-CD4-GFP

To insert a *UAS-CD4-GFP* transgene at the *attP40* equivalent site of *D. sechellia*, we integrated a *UAS-CD4-GFP* cassette^83^ into *Dsec-attP40-3xP3-DsRed* via Gibson assembly.

#### Flp-out constructs

We amplified the flp-out cassette from *pJFRC208-10XUAS-FRT>STOP>FRT-myr::smGFP-FLAG*^53^ (Addgene #63169) via PCR and integrated it into *Dsec1029-mhc-YFP* via Gibson assembly resulting in *Dsec1029-flip-out-mhc-YFP*. Similarly, *DsecattP40-flip-out-3xP3-DsRed* was generated by PCR amplification of *pJFRC208-10XUAS-FRT>STOP>FRT-myr::smGFP-HA*^53^ (Addgene #63166) and integration into *Dsec-attP40-3xP3-DsRed*.

#### Single guide RNA expression vectors

For expression of individual short guide (sg)RNAs, oligonucleotide pairs (**Sup. Table 6**) were annealed and cloned into BbsI-digested *pCFD3-dU6-3gRNA* (Addgene #49410), as previously described^84^. To express multiple sgRNAs from the same vector backbone, oligonucleotide pairs (**Sup. Table 7**) were used for PCR and inserted into *pCFD5* (Addgene #73914) via Gibson Assembly, as previously described^85^.

### Insertion site mapping

Mapping of piggyBac insertions using TagMap was performed as previously described^86^. Sequencing hits were blasted against the *D. sechellia* ASM438219v2 genome assembly^87^. Locations of transgene insertion sites are listed in **Sup. Table 1**.

### Flp-out mediated stochastic labelling

For stochastic labelling, we combined *D. sechellia* Gal4 driver lines with *Dsec-hs-Flp1.* The resulting double-transgenic lines were crossed to *Dsec-attP1029-flip-out-mhc-YFP*, *DsecattP40-flip-out-3xP3-DsRed* or both. In *D. melanogaster*, we combined *D. melanogaster* Gal4 driver with MCF0-1^53^. F1 progeny was heat shocked for 3-5min at 37°C in an incubator at the larval L3 stage, 24 h, 48 h, or 72 h after pupa formation. Brains of emerging flies were dissected and stained with combinations of anti-GFP, anti-FLAG, anti-HA, anti-Cadherin and anti-nc82 antibodies as described in the immunohistochemistry section.

### Immunohistochemistry

Immunofluorescence on adult brains was performed as previously described^88^ using mouse monoclonal antibody nc82 1:5, rat anti-Cadherin 1:25 (both Developmental Studies Hybridoma Bank), rat anti-HA 1:500 (Roche), rabbit anti-FLAG 1:500 (Novus Biologicals), and rabbit anti-GFP 1:500 (Invitrogen). Alexa488-, Cy3- and Cy5-conjugated goat anti-mouse, goat anti-rat and goat anti-rabbit IgG secondary antibodies (Molecular Probes, Jackson Immunoresearch) were used at 1:500.

### Image acquisition and processing

Confocal images of brains were acquired on an inverted confocal microscope (Zeiss LSM 710) equipped with an oil immersion 40x objective (Plan Neofluar 40ξ oil immersion DIC objective; 1.3 NA), unless stated otherwise. For whole brain z-stacks, a x/y dimensions of 768ξ768px (353,8ξ353,8µm) was used and slice thickness was set to 1µm without slice overlap. The pinhole was set to 1AU (∼2.17µm). The number of slices ranged between 170-190. Images were processed in Fiji^89^. *D. sechellia* brains were registered to the *D. sechellia* reference DsecF and subsequently to DsecI^29^ and *D. melanogaster* brains to the IS2 *D. melanogaster* reference brain using the Fiji CMTK plugin (https://github.com/jefferis/fiji-cmtk-gui), as previously described^90^.

#### Antennal lobe segmentation & neuron reconstruction

Segmentation of the *Dsec* reference brain AL was performed using Amira 6.5 (Thermo Fisher Scientific) based on similar re-constructions^34,56^. Projection neuron (PN) morphologies were manually traced using neuTube 1.0z^91^. PN identity was assigned based on dendritic glomerular innervation in the AL, axonal branching pattern in the LH, and NBLAST clustering.

#### Comparison of neuron morphologies

Intra- and interspecific comparisons of PNs was performed using the R-based natverse ecosystem^92^. For *D. melanogaster*, we extracted neuron traces of adult AL projection neurons generated by light imaging and connectomic reconstructions from the Virtual Fly Brain collection^57^ (search tag: “adult antennal lobe projection neuron”, filtering for confocal neurons, FAFB, and selection for adPN or lPN types) and bridged them from the JRC2018U to the IS2 reference space^90^. Traces were not re-rooted (resulting in some erroneous annotations of cell soma). The dataset curation is described in detail in the provided R scripts. *D. sechellia* traces were bridged from DsecI to IS2 and all left-hemisphere neurons were mirrored to the right hemisphere. For intraspecific analysis, *D. sechellia* neurons were bridged to DsecI. (Traces are depicted in orthogonal view).

#### Measurements of branch length

Branch length measurements were performed on traced neurites using the simple neurite tracer toolbox^93^.

#### Connectomic circuit identification

Potential downstream neurons of *D. sechellia* PN innervations where identified in the *D. melanogaster* hemibrain (v1.2.1) by spatial proximity using neuprint^94^.

### *In vivo* two-photon calcium imaging and photoactivation

Two-photon calcium imaging, odour delivery, image analysis and photoactivation was performed as described previously^33^ with the exception that for odour delivery, the constant air flow in the olfactometer was not humidified. Noni juice (Raab Vitalfood) was diluted in water. Methyl hexanoate (CAS 106-70-7) and *2*-heptanone (CAS 110-43-0) were purchased in the highest purity from Sigma Aldrich and diluted in dichloromethane (Sigma Aldrich, CAS 75-09-2).

### Statistics and reproducibility

Data were analysed and plotted using Excel and R (v3.2.3; R Foundation for Statistical Computing, Vienna, Austria, 2005; R-project-org).

### Data, code and biological material availability

All relevant data supporting the findings of this study, code used for analysis and all unique biological materials generated in this study (e.g., mutants, plasmids and transgenic fly strains) are available from the corresponding author upon request. Neuron tracings and code are available at the following github repository: https://github.com/AuerTomLab/Duerr2024. Tracings are also included in Supplementary Data.

---

**Supplementary Figure 1. *attP* landing site establishment in *D. sechellia*.**

**a)** Left: Schematic of the insertion cassettes used for piggyBac- and CRISPR-Cas9-mediated integration. The *FRT* and *mFRT71* sites placed between promoter and fluorescent marker do not interfere with expression. MHC = *myosin heavy chain* promoter (673bp version^82^). Right: Integration sites established with these markers at random (piggyBac) and homologous locations to characterised sites in *D. melanogaster* (*attP2*^45^, *attP40*^42^, *JK22C*^44^, *su(Hw)attP2*^95^) and *D. simulans* (insertion no.1029^47^).

**b)** Location of *attP* sites on the four chromosomes of the *D. sechellia* genome. Black dot = centromere. Indicated below selected sites: transgenic constructs integrated via PhiC31-mediated transgenesis and used in panels **c,d** and **Fig. 1**. Some transgenes have been published in Ref.^33^ as indicated.

**c)** Labelling of the DM2, DA1, and DL1 glomeruli by a split-Gal4 combination in *D. sechellia* combined with a *20xUAS-GCaMP6s* reporter. The 2-3 additional glomeruli shown in **Fig. 1d** are not visible with this reporter – expression in DM2 (black arrow) is weak. Scale bar = 50 µm. Insets: labelling of DM2. Scale bar = 5 µm.

**d)** Anti-GFP immunofluorescence of the *D. sechellia* central brain of two *UAS*-reporter lines without Gal4 driver. The reporter integrated at the *Dsec-attP40* locus shows weak GFP expression in Kenyon cells (black arrows). Scale bar = 50 µm.

**Supplementary Figure 2. Overview of *D. sechellia* PN cell types.**

**a)** Top: schematics of the Flp-cassette at the *Dsec-attP40* locus. Below: schematics of the Flp-cassette at the *Dsec-attP1029* locus and potential outcomes of Flp recombinase-mediated recombination events. The *attP* landing site marker contains already a *FRT* site, resulting in multiple possible recombination events. Only outcome 3 leads to functional reporter expression (further increasing label sparseness with this transgene).

**b)** Classification of the 93 traced uPNs into 33 uniPNs and 60 uni^+^PNs.

**c)** Overview of the 26 PN types identified in the *D. sechellia* dataset. Left hemisphere: all neurons of the respective type; right hemisphere: individual representative neuron. Blue: segmentation of AL glomerulus/i with main dendritic innervation. Top row: 6 types of mPNs. 2 middle rows: uPNs with more than 3 examples in the dataset. Bottom row: uPNs with < 3 examples in the dataset.

**Supplementary Figure 3. Conserved PN morphologies between *D. melanogaster* and *D. sechellia*.**

**a)** Comparison of mlPN1 morphologies between *D. sechellia* (left, light microscopy, LM) and *D. melanogaster* (right, electron microscopy, EM). Here, no individual examples are shown as axons of neurons with soma in the left hemisphere cross to the right hemisphere. The shorter contra-lateral branch in the *D. melanogaster* EM example could result from incomplete tracing across hemispheres.

**b)** Dorsal madPN1s in *D. sechellia* and *D. melanogaster*.

**c)** Comparison of seven uPN types between *D. sechellia* (left, LM) and *D. melanogaster* (right, LM and EM) with conserved neuroanatomy.

**Supplementary Figure 4. Specific labelling of PN subtypes via photoactivation.**

**a)** Photoactivation of the antero-medial branch area in *D. sechellia* expressing photo-activatable GFP in most PNs results in labelling of the DM2 and VM5d glomeruli. Left: Picture of the whole brain with labelling of the branch area and DM2 and VM5d innervating dendrites in the AL. Scale bar = 50 µm. Right: higher magnification pictures of the AL showing DM2 (top) and VM5d (bottom) innervations.

**b)** Photoactivation of VM5d innervating PNs in *D. simulans* expressing photo-activatable GFP pan-neuronally. Left: Picture of the whole brain with labelling of VM5d and projections to the LH. Scale bar = 50 µm. Right, top: Higher magnification pictures of the LH. White arrowheads: medial AL tract (mALT), medial inhibitory AL tract (mIALT), and inhibitory AL tract (IALT). No branching into the antero-medial brain region is detected. Bottom: Overview of the AL with only the VM5d glomerulus being labelled. Scale bar = 10 µm.

**Supplementary Figure 5. Potential *D. sechellia* VM5d/DM2 PN downstream partners based on connectomic data in *D. melanogaster*.**

**a)** Same as **Fig. 3j**: relative location of the *D. sechellia* specific axonal PN branch between the superior lateral protocerebrum (SLP) and the superior clamp (SCL). White line anterior view: plane of cross-section panel **b**; white line dorsal view: plane of cross-section panel **c**.

**b,c)** EM section along (**b**) and perpendicular (**c**) to the axonal tract region. Red borders: novel PN innervations in *D. sechellia*. Blue: axonal tract. Several neurons (in colours) in *D. melanogaster* are in close proximity and potential downstream partners in *D. sechellia*. For a complete list of neurons in this target area, see **Sup. Table 4**.

**d)** 3D reconstruction of neurons indicated in **b, c** in *D. melanogaster* together with VM5d PNs of *D. sechellia* (red, in the IS2 reference space). Either individual neurons (top) or neuron classes (bottom) are shown. Black dots = neuron somata.

