## Supplementary figures and images for "Olfactory projection neuron rewiring in the brain of an ecological specialist"

### Supplementary Figures

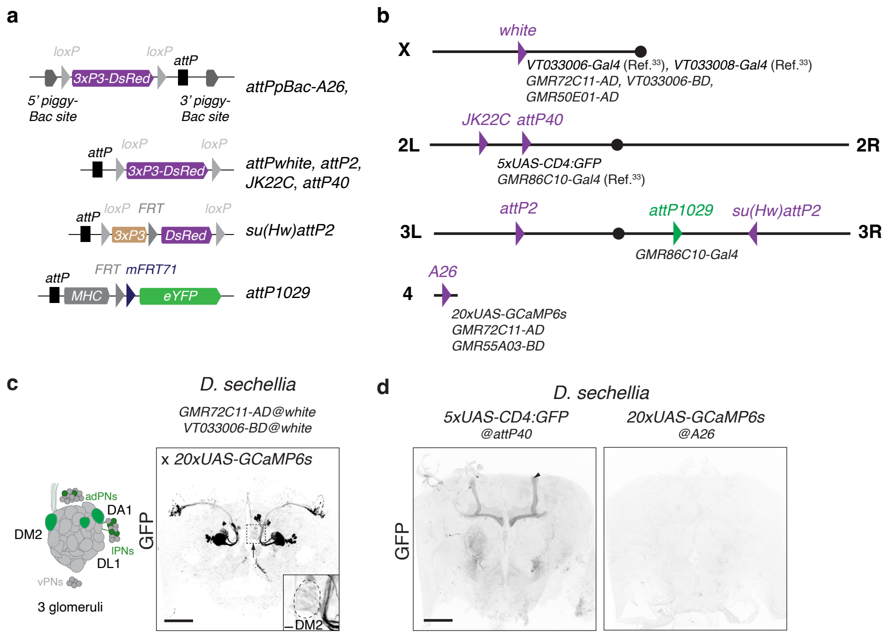

#### Supplementary Figure 2

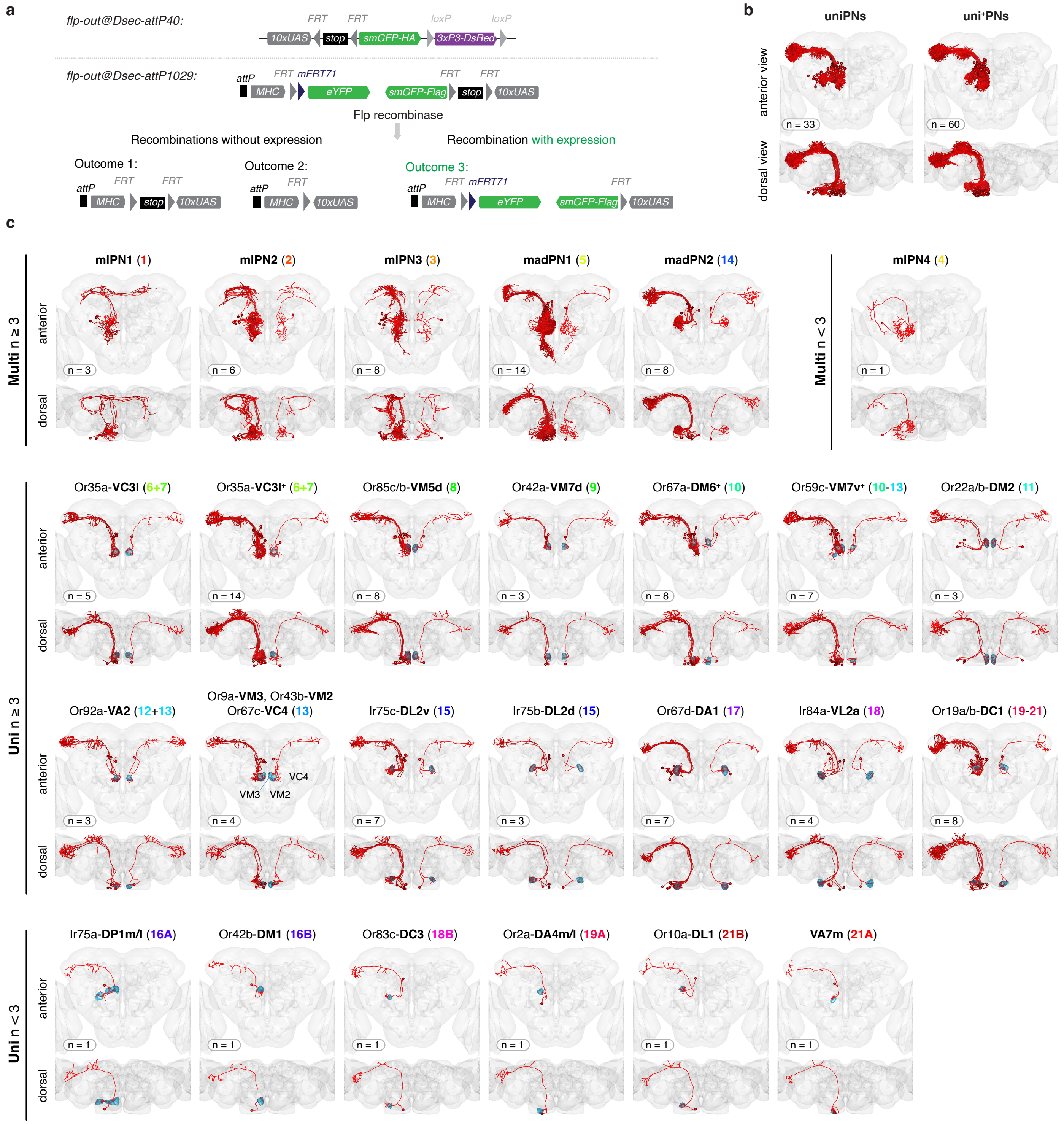

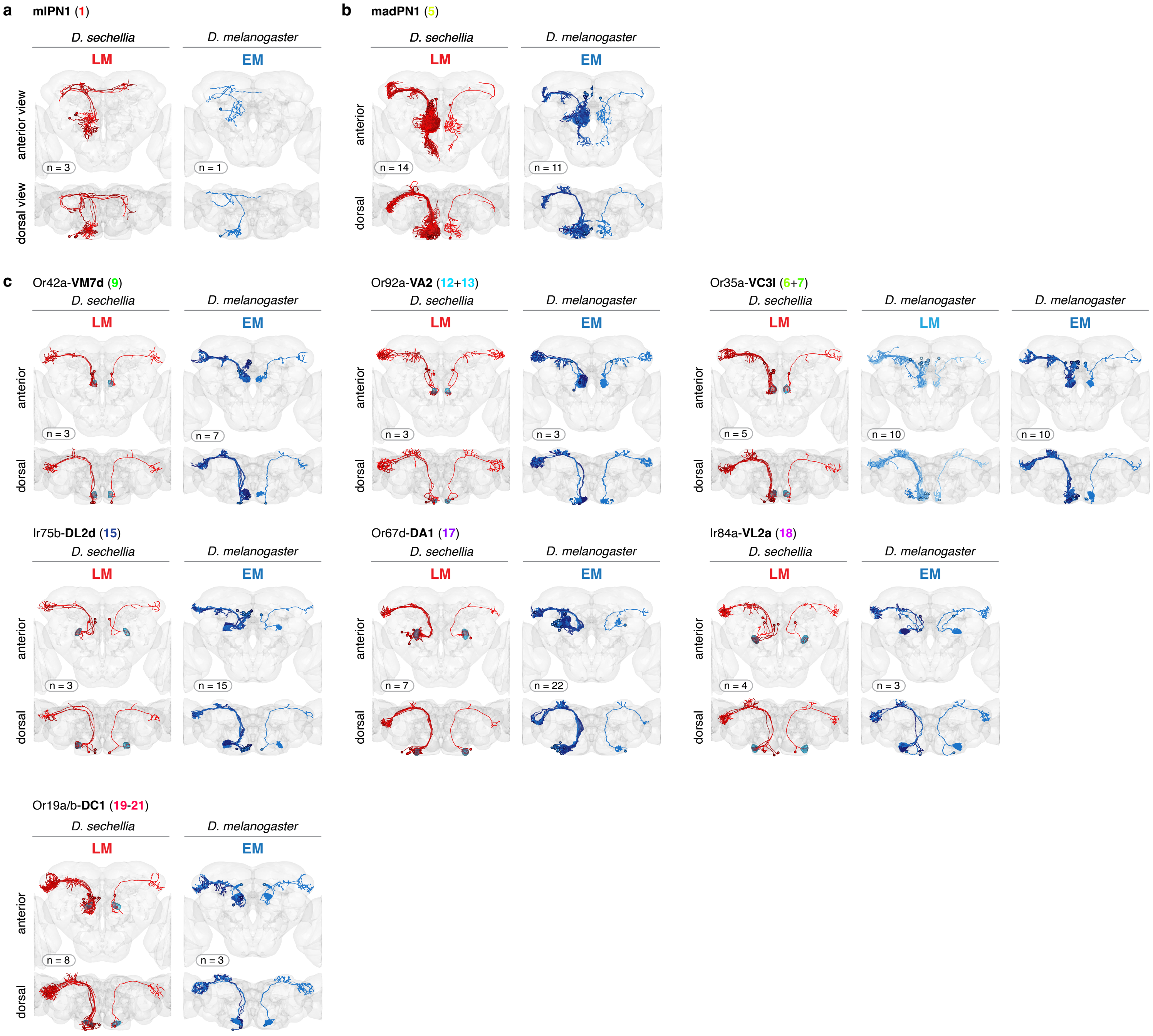

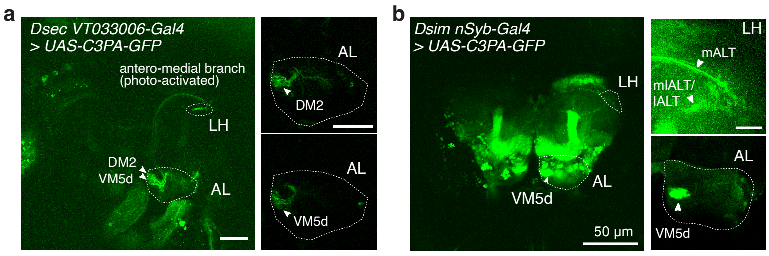

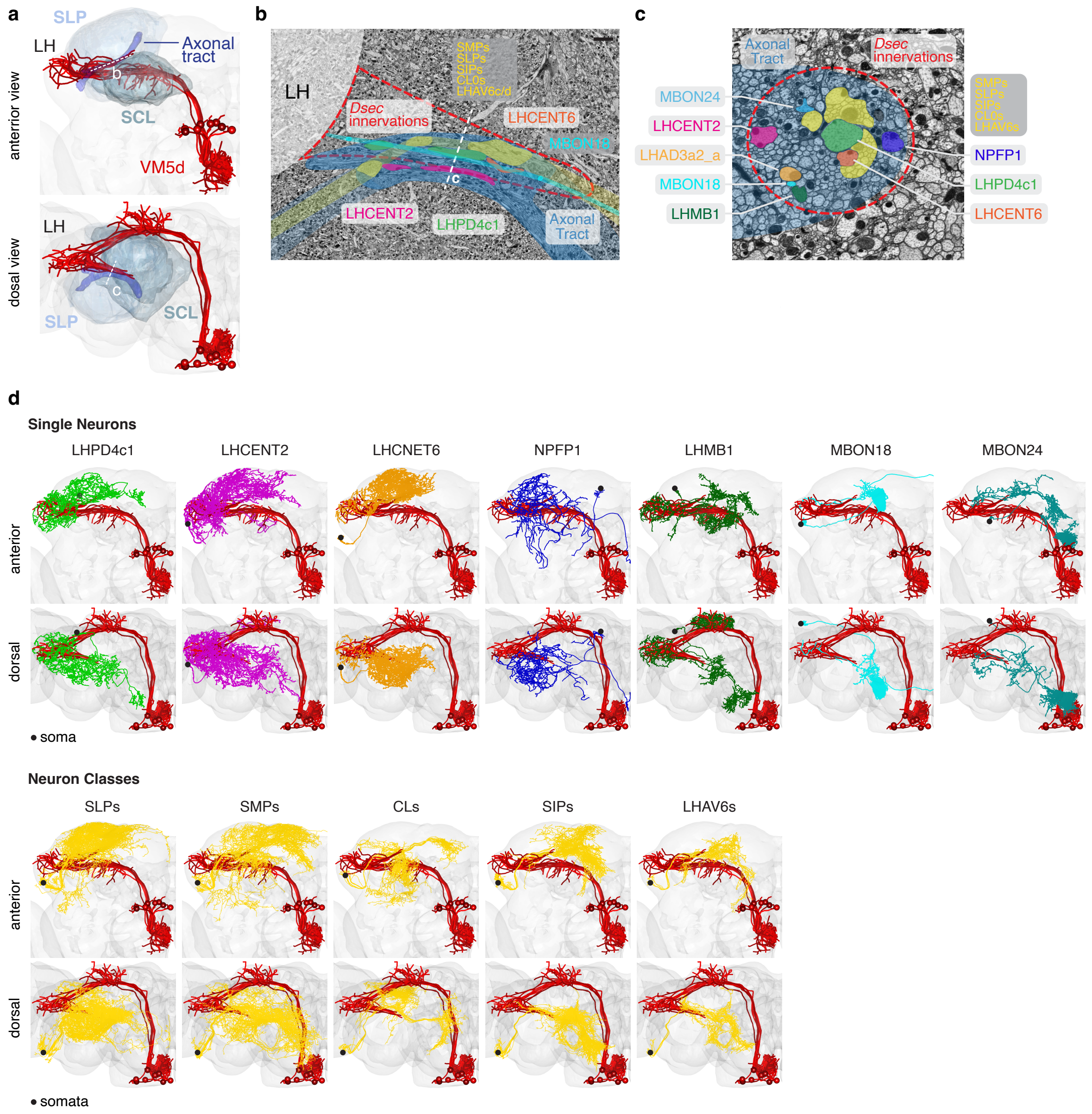
