## Supplementary Tables 2,5,6,7 for "Olfactory projection neuron rewiring in the brain of an ecological specialist"

**Supplementary Table 1.** Genomic locations of piggyBac insertions. In blue = *attP* site used in this study.

**Supplementary Table 2:** Efficiencies of insertions at *attP* locations

| ***attP* site tested** | **Construct** | **No. embryos injected** | **No. fertile crosses** | **No. transgenic progeny** | **Efficiency (transgenic progeny/fertile crosses)** |
|---|---|---|---|---|---|
| *Dsec-attP1029* | *GMR_86C10Gal4* | 600 | 98 | 1 | 1% |
| *Dsec-attP40* | | 762 | 77 | 1 | 1.3% |
| *Dsec-attP2* | | 940 | 76 | 0 | 0 |
| *Dsec-attPpBac-A26* | | 525 | 56 | 0 | 0 |
| *Dsec-attPsu(Hw)attP2* | | 560 | 51 | 0 | 0 |
| *Dsec-attPJK22C* | | Homozygous lethal in *D. sechellia* | | | |
| *Dsec-attPpBac-A26* | *p(GP-JFRC7-20XUAS-IVS-GCaMP6s)* | 355 | 41 | 1 | 2.4% |
| *Dsec-attP40 locus/sgRNAs* | CRISPR KI *UAS-CD4-GFP* | 240 | 37 | 1 | 2.7% |

Integration efficiencies (especially at *Dsec-attP2* and *Dsec-attPsu(Hw)attP2*) might be higher when aiming for integration of smaller constructs. For CRISPR-mediated integration at the *attP40* locus, a sgRNA expression vector *pCFD5-Dsec-attP40* and *Dsec-attP40-3xP3-DsRed-UAS-CD4:GFP* were co-injected.

**Supplementary Table 3.** List of all reconstructed PNs in *D. sechellia*. ID = neuron identity (see Supplementary Data for swc tracing files); PN type: adPN = anterior-dorsal PN cluster, lPN = lateral PN cluster; branching type: up = uni+-glomerular branching, u = uni-glomerular branching, m = multi-glomerular branching; glomerulus innervation = main target glomerulus, additional innervation = additional target glomeruli; hemisphere: R = right, L = left; driver = genetic Gal4 driver used for stochastic labelling.

**Supplementary Table 4.** List of potential PN downstream neurons based on EM circuit data.

**Supplementary Table 5.** Wildtype and transgenic lines used and generated in this study.

| **Application** | **Stock name** | **Donor plasmid** | **Parental strain** | **Species** | **Method/Reference** | **Figure** |
|---|---|---|---|---|---|---|
| *wildtype* | *Dsec.07 (wildtype)* | | *Drosophila* Species Stock Center [DSSC] 14021-0248.07 | *D. sechellia* | | |
| | *Dsec.30 (white mutant)* | | DSSC 14021-0248.30 | *D. sechellia* | | |
| *attP/B transgenesi* | *Dsec-attPpBac-A26* | *pBac-attP-3P3-DsRed* | DSSC 14021- | *D. sechellia* | piggyBac integration this | Fig. 1 |

| | | | | | | |
|---|---|---|---|---|---|---|
| *s* | | | 0248.07 | | study | |
| | *Dsec-attP40* | *pHD-DsecattP40-3P3-DsRed* | *Dsecnanos-Cas9*[29] | *D. sechellia* | CRISPR knock-in, this study | Fig. 1 |
| | *Dsec-attP2* | *pHD-DsecattP2-3P3-DsRed* | *Dsecnanos-Cas9*[29] | *D. sechellia* | CRISPR knock-in, this study | Sup. Fig. 1 |
| | *Dsec-attPwhite* | | DSSC 14021-0248.07 | *D. sechellia* | [29] | Fig. 1 |
| | *Dsec-attP1029* | *pHD-Dsecatt1029-MHC-YFP* | *Dsecnanos-Cas9*[29] | *D. sechellia* | CRISPR knock-in, this study | Fig. 1 |
| | *Dsec-attPJK22C* | *pHD-DsecJK22C-3P3-DsRed* | *Dsecnanos-Cas9*[29] | *D. sechellia* | CRISPR knock-in, this study | Sup. Fig. 1 |
| | *Dsec-su(Hw)attP2* | *pHD-Dsecsu(Hw)attP2-3P3-DsRed* | *Dsecnanos-Cas9*[29] | *D. sechellia* | CRISPR knock-in, this study | Sup. Fig. 1 |
| *Gal4-driver* | *Dmel-GMR86C10-Gal4* | | Bloomington *Drosophila* Stock Center [BDSC] 46820 | *D. melanogaster* | | Fig. 1, Fig. 4, Fig. 5 |
| | *Dmel-VT033006-Gal4* | | BDSC_73333 | *D. melanogaster* | [96] | Fig. 1, Fig. 5, |
| | *Dsec-GMR86C10-Gal4@Dsec-attP40* | *Dmel GMR_86C10-Gal4* | *Dsec-attP40* | *D. sechellia* | [33] | Fig. 1, Fig. 4, Fig. 5 |
| | *Dsec-GMR86C10-Gal4@Dsec-attP1029* | *Dmel GMR_86C10-Gal4* | *Dsec-attP1029* | *D. sechellia* | attB/P integration, this study | Fig. 1 |
| | *Dsec-VT033006-Gal4* | | | *D. sechellia* | [33] | Fig. 1, Fig. 2, Fig. 5, Sup. Fig. 4 |
| | *Dsec-VT033008-Gal4* | | | *D. sechellia* | [33] | Fig. 2 |
| | *Dmel-SS0186F* | *Gift from G. Jefferis* | SS0186F Split Gal4 line | *D. melanogaster* | [24] | Sup. Fig. 1 |
| | *Dsec-VT033006 pGag4BD@white* | *VT033006 pGAL4BD (Gift from G. Rubin)* | *Dsec-attPwhite*[29] | *D. sechellia* | attB/P integration, this study | Fig. 1, Sup. Fig. 1 |
| | *Dsec-GMR72C11-p65AD@white* | *GMR72C11-p65AD (Gift from G. Rubin)* | *Dsec-attPwhite*[29] | *D. sechellia* | attB/P integration, this study | Fig. 1, Sup. Fig. 1 |
| | *Dsec-GMR72C11-p65AD@Dsec-attPA26* | *GMR72C11-p65AD (Gift from G. Rubin)* | *Dsec-attPpBac-A26* | *D. sechellia* | attB/P integration, this study | Fig. 1 |
| | *Dmel-SS00587* | *Gift from G. Jefferis* | SS00587 Split Gal4 line | *D. melanogaster* | [52] | Sup. Fig. 1 |
| | *Dsec-GMR55A03-pGal4DB@Dsec-attPA26* | *GMR55A03-pGAL4DB (Gift from G. Rubin)* | *Dsec-attPwhite*[29] | *D. sechellia* | attB/P integration, this study | Sup. Fig. 1 |
| | *Dsec-GMR50E01-p65AD@white* | *GMR50E01-p65AD (Gift from G. Rubin)* | *Dsec-attPpBac-A26* | *D. sechellia* | attB/P integration, this study | Sup. Fig. 1 |
| | *Dsim-VT033006Gal4* | *pVT033006-Gal4*[96] | *Dsim1029* | *D. simulans* | attB/P integration, this study | Fig. 5 |
| | *DsimNsyb-Gal4* | Gift from S. Caron | | *D. simulans* | [34] | Sup. Fig. 4 |
| *UAS-* | *Dsec-20xUAS-* | *p(GP-JFRC7-* | *Dsec-* | *D. sechellia* | attB/P integration, | Fig. 4, |

| | | | | | | |
|---|---|---|---|---|---|---|
| *reporter* | *GCamP6s* | *20XUAS-IVS-GCaMP6s)*[97] | *attPpBac-A26* | | this study | Fig. 5, Sup. Fig. 1 |
| | *Dsec-UAS-CD4:GFP* | | *Dsec*nanos-Cas9[29] | *D. sechellia* | CRISPR knock-in, this study | Fig. 1, Sup. Fig. 1 |
| | *Dsec-UAS-C3PA-GFP* | | | *D. sechellia* | [29] | Sup. Fig. 4 |
| | *Dsec-UAS-BrpStraw* | *pUAS-BrpStraw (Gift from S. Sigrist)*[61] | | *D. sechellia* | P-element integration, this study | Fig. 5 |
| | *Dsec-UAS-GCamP6f* | | | *D. sechellia* | [29] | Fig. 5 |
| | *Dmel-5xUAS-mCD8:GFP* | | | *D. melanogaster* | [98] | Fig. 1 |
| | *Dmel-20xUAS-GFP* | | BDSC_52262 | *D. melanogaster* | | Fig. 1 |
| | *Dmel-20xUAS-GCaMP6s* | | BDSC_ 42746 | *D. melanogaster* | | Fig. 4, Fig. 5 |
| | *Dsim-UAS-GCaMP6s* | | | *D. simulans* | [60] | Fig. 5 |
| | *Dsim-UAS-C3PA-GFP* | | | *D. simulans* | [34] | Sup. Fig. 4 |
| *Stochastic labelling* | *Dsec-flp-out@Dsec-attP40* | | *Dsecnanos-Cas9*[29] | *D. sechellia* | CRISPR knock-in, this study | Fig. 2 |
| | *Dsec-flp-out@Dsec-attP1029* | | *Dsecnanos-Cas9*[29] | *D. sechellia* | CRISPR knock-in, this study | Fig. 2, Fig. 4 |
| | *Dsec-hs-Flp1* | *pBPhsFlp1 (Addgene #32148)* | *Dsec-attPpBac-A26* | *D. sechellia* | attB/P integration, this study | Fig. 2, Fig. 4 |
| | *Dmel*-MCF0-1 | | BDSC_64085 | *D. melanogaster* | [53] | Fig. 4 |

**Supplementary Table 6.** Oligonucleotides used to generate single sgRNA expression vectors.

| **Target** | **Forward primer (5'-3')** | **Reverse primer (5'-3')** |
|---|---|---|
| *Dsec-attPJK22C sgRNA1* | GTCGCATCAGCAACGCACAGTGGG | AAACCCCACTGTGCGTTGCTGATG |
| *Dsec-attPJK22C sgRNA2* | GTCGACTGTGCGTTGCTGATGCAG | AAACCTGCATCAGCAACGCACAGT |
| *Dsec-attP2 sgRNA1* | GTCGGCGAATGGCGAAAGCTGCGC | AAACGCGCAGCTTTCGCCATTCGC |

**Supplementary Table 7.** Oligonucleotides used to generate multi-sgRNA expression vectors.

| **Target** | **Name** | **Sequence (5'-3')** | **Resulting sgRNAs** |
|---|---|---|---|
| *Dsec-su(Hw)attP2* | *PCR1fwd* | GCGGCCCGGGTTCGATTCCCGGCCGATGCAGGAATCGCACGGACGTGTCGGTTTTAGAGCTAGAAATAGCAAG | sgRNA1: GGAATCGCACGGACGTGTCG |
| | *PCR1rev* | ATTTTAACTTGCTATTTCTAGCTCTAAAACGCAGATTCTCACTTCATGCTTGCACCAGCCGGGAATCGAACCC | sgRNA2: AGCATGAAGTGAGAATCTGC |
| *Dsec-attP40* | *PCR1fwd* | GCGGCCCGGGTTCGATTCCCGGCCGATGCAGTTACACAGCCCTTGGCTGGGTTTTAGAGCTAGAAATAGCAAG | sgRNA1: GTTACACAGCCCTTGGCTGG |

| | | | |
|---|---|---|---|
| | *PCR1rev* | ATTTTAACTTGCTATTTCTAGCTCTAAAACACACACATTGG GCAGCACAGTGCACCAGCCGGGAATCGAACCC | sgRNA2: CTGTGCTGCCCAATGTGTGT |
| *Dsec-attP1029* | *PCR1fwd* | GCGGCCCGGGTTCGATTCCCGGCCGATGCATTAAAGTTA AACAAGCGGTAGTTTTAGAGCTAGAAATAGCAAG | sgRNA1: TTAAAGTTAAACAAGCGGTA |
| | *PCR1rev* | ATTTTAACTTGCTATTTCTAGCTCTAAAACTTTCTACTTTCA GATTGGTTTGCACCAGCCGGGAATCGAACCC | sgRNA2: AACCAATCTGAAAGTAGAAA |
